# Gamma Approximation of Stratified Truncated Exact test (GASTE-test) & Application

**DOI:** 10.1101/2024.07.26.605317

**Authors:** Alexandre Wendling, Clovis Galiez

## Abstract

The analysis of binary outcomes and features, such as the effect of vaccination on health, often rely on 2 × 2 contingency tables. However, confounding factors such as age or gender call for stratified analysis, by creating sub-tables, which is common in bioscience, epidemiological, and social research, as well as in meta-analyses. Traditional methods for testing associations across strata, such as the Cochran-Mantel-Haenszel (CMH) test, struggle with small sample sizes and heterogeneity of effects between strata. Exact tests can address these issues, but are computationally expensive. To address these challenges, the Gamma Approximation of Stratified Truncated Exact (GASTE) test is proposed. It approximates the exact statistic of the combination of p-values with discrete support, leveraging the gamma distribution to approximate the distribution of the test statistic under stratification, providing fast and accurate p-value calculations, even when effects vary between strata. The GASTE method maintains high statistical power and low type I error rates, outperforming traditional methods by offering more sensitive and reliable detection. It is computationally efficient and broadens the applicability of exact tests in research fields with stratified binary data. The GASTE method is demonstrated through two applications: an ecological study of Alpine plant associations and a 1973 case study on admissions at the University of California, Berkeley. The GASTE method offers substantial improvements over traditional approaches and is available as an open-source package at https://github.com/AlexandreWen/gaste.

## 1. Introduction

Categorical data analysis is ubiquitous across various scientific fields, particularly in examining binary outcomes in relation to binary features, such as the influence of a vaccination on individuals’ health. The objective often involves testing whether the probability of outcomes (sick or healthy) differs between two groups (vaccinated or not). To investigate the association between these features and outcomes, a 2 × 2 contingency table is frequently used to summarize binary counts within a sample. However, observations may be influenced by external factors. To address this, stratification is utilized to create sub-tables based on these confounding variables, such as age or gender. This multilevel analysis approach is frequently used in bioscience studies, epidemiology, social research, and meta-analyses (Agresti, 2012; Smaardijk et al., 2020; Reilly, 2023).

To test the overall association between binary features and outcomes across several strata, one way is to calculate a combined effect given by the weighted average of some summary statistic across all studies. Two models can be considered for assigning weights (Borenstein et al., 2010), a fixed-effects model assumes that all strata estimate the same underlying effect size (the strength of association), whereas a random-effects model accounts for variability both within and between studies. The latter assumes that the true effect size varies among studies, and the observed effect sizes reflect this distribution. Four methods are widely used, three fixed effects methods, namely Mantel-Haenszel (Mantel and Haenszel, 1959), Peto (Yusuf et al., 1985) and inverse variance; and one random effects method, DerSimonian and Laird inverse variance (DerSimonian and Laird, 1986). These methods employ asymptotic tests that are hampered by small counts and do not handle heterogeneity (uneven strength of association between feature and outcome across strata) even in random-effects models due to averaging over strata. Chapter 10 of the Cochrane Handbook provides useful insights and a clear summary of the topic (Deeks JJ, 2023). The indispensable stratification carried out in studies to manage confounding variables increases the number of combinations of subgroup categories and creates problems known as *combination fatigue*, including concerns about small sample size, reduced power, spurious results and complexity in statistical design and analysis (Evans, 2024). However, as we have mentioned, despite their widespread use, the aforementioned methods may lack robustness and power in these configurations. This is particularly true for the Mantel-Haenszel test (Mantel and Haenszel, 1959) also called the Cochran-Mantel-Haenszel (CMH) test (Cochran, 1954), despite its very popular use, which is sometimes misapplied (Berger et al., 2006).

Leveraging many strata without being constrained by rare events (low cell frequency) or heterogeneity of association requires exact testing. In essence, an exact test aims to determine the probability of observing a specific stratified set of tables under a null model stating that the outcomes (e.g., sick or healthy) are independent of the features (vaccinated or not). Given the number of samples in each table, there exists a finite number of possibilities for different stratified tables, allowing the calculation of the exact probability of the observed data. No assumptions of homogeneity or sufficient counts are henceforth needed. Exact tests based on this principle have already been proposed (Zelen, 1971; Agresti, 1992; Jung, 2014; Chiba, 2017), in particular by considering as the test statistic the sum, across strata, of the positive outcomes having the feature. This way of building the statistic makes the power highly dependent on homogeneity of the effect size across strata, as a small count in one stratum can compensate for a large one in another. To mitigate this effect, we propose a different approach: adapting the Fisher’s combination (sum of log of uniformly distributed *p*-values) to the case of sub-uniform *p*-values obtained from hypergeometric one-tailed tests, at each stratum.

Despite the great interest in exact tests, their computational complexity grows exponentially with the number of strata and the number of samples in each stratum, making the test intractable in many scenarios. To address the complexity issue, drawing inspiration from the combination of dependent *p*-values (Brown, 1975; Kost and McDermott, 2002; Yang, 2010; Zheng et al., 2012; Li et al., 2014), we propose an approximation of this *p*-value combination using a gamma distribution. We infer the gamma distribution parameters by matching its moments with the analytical moments of the exact distribution. This approach is particularly encouraged by the recent work of Contador and Wu (2023) demonstrating that the probability distribution within the Gamma family that minimizes the Wasserstein distance to the exact distribution of the statistic of the *p*values combination is given by the parameters matching the two first moments. We will discuss the relevance of the choice of the Wasserstein distance in the context of statistical testing. The method of moment matching is applied to the truncated case, where the combination statistic takes only into account the *p*-values of strata that pass a user-defined significance threshold, to increase the significance of the test in case of heterogeneity of the effect across strata. In contrast to asymptotic approximations, directly approximating the exact test allows handling cases of rare events, particularly in the case of a high number of strata, leading to a high complexity of exact calculation. We call our test method GASTE-test for Gamma Approximation of Stratified Truncated Exact test. Moreover, this approximation allows us to reduce the computational complexity by more than three orders of magnitude and makes the exact test, by its approximation, more practical for real-world applications. We provide simulations to show the actual control of the type I error rate and show that the GASTE test is, on average, 16% more powerful than the CMH test when the effect is applied to all strata and 58% more powerful when the effect is applied to only half the strata while controlling the type I error level.

We also illustrate the efficacy of our method through real-world applications, comparing it to other statistical tests to assess the overall significance between outcomes and features in stratified 2 × 2 contingency tables. The initial application of our test was in the field of ecology, which spurred its development. We had multiple observation zones in the French Alps, and within each zone, points were recorded for the presence or absence of various plant species. For a given pair of plants, we constructed stratified tables by environment to study plant associations by counting coexistence. This application was particularly interesting because the number of survey points was relatively low (≤300) and was not controlled, and within these observation zones, species could be either rare or very abundant. The GASTE-test method stands out from the rest by detecting twice as many associations as the CMH test, and handles the heterogeneity of effects between strata, which is crucial in this context. The second application is a well-known case study of stratified binary data. It puts at stake whether the admissions to the University of California at Berkeley in 1973 are gender biased. When aggregated, the data appear to show a selection bias favoring men. However, when stratified by department, the usual tests give information on the heterogeneity of the association between strata and surprisingly no overall association between gender and department. The GASTE-test method gives a different result.

## 2. Methods

### 2.1. Stratified 2×2 contingency table

When dealing with observations of individuals with binary features (e.g., vaccinated or not), associated with binary outcomes (e.g., sick or healthy), it is convenient to represent the data as a 2 × 2 contingency table. Observations may be influenced by some external factors, to control these confounding variables, stratification can be used to create sub-tables based on these factors. Assuming S strata *s* = (1, …, *S*), Table 1 for a given stratum *s* contains the following data: *k*_*s*_ number of positive outcomes with the feature, i.e., value of co-occurrence, *K*_*s*_ total number of positive outcomes, *n*_*s*_ total number of features and *N*_*s*_ the sample size.

**Table 1:** 2×2 contingency table for stratum *s*

| Outcomes | Features |  | Total |
| --- | --- | --- | --- |
|  | Yes | No |  |
| Positive | $k_s$ | $K_s - k_s$ | $K_s$ |
| Negative | $n_s - k_s$ | $N_s - n_s - K_s + k_s$ | $N_s - K_s$ |
| Total | $n_s$ | $N_s - n_s$ | $N_s$ |

**Table 2:** Family-wise error rate (FWER), under the null hypothesis, for the different association detection methods on ORCHAMP data.

|  |  | GASTE | Bonferroni | TFisher | CMH |
| --- | --- | --- | --- | --- | --- |
| $\alpha = 0.2$ | under-assoc | 0.0475 | 0.0125 | 0.0125 | 0.015 |
|  | over-assoc | 0.05 | 0.015 | 0.015 | 1 |
| $\alpha = 0.05$ | under-assoc | 0.015 | 0.005 | 0.0025 | 0.005 |
|  | over-assoc | 0.015 | 0.005 | 0.005 | 1 |
| $\alpha = 0.01$ | under-assoc | 0.005 | 0.0 | 0.0 | 0.0 |
|  | over-assoc | 0.005 | 0.0025 | 0.0025 | 1 |

### 2.2. Fisher exact test

For a stratum *s*, the ratio 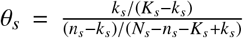, called the odds ratio, is used to test the association between features and outcomes. When the outcome is independent of the features, the ratio *θ*_*s*_ is close to 1. Indeed, if the outcomes are not affected by the presence or absence of features, i.e., if a positive outcome appears randomly with or without the feature, the ratio of positive outcomes in the case of feature to those in the case of a non-feature must be similar to the ratio of negative outcomes between feature and non-features.

So in this case, with fixed marginals *n*_*s*_, *K*_*s*_ and fixed sample size *N*_*s*_ (conditional test), the random variable *X*_*s*_ representing the number of individuals with both a positive feature and a positive outcome follows a hypergeometric distribution (*HG*). Formally, the probability of observing *k*_*s*_ positive outcomes with the feature is given by 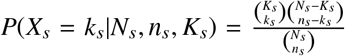, that is, also the exact probability of observing a specific table with fixed marginals. The support of the distribution is discrete and finite and is given by supp(*X*_*s*_) = ⟦max(0, *n*_*s*_ +*K*_*s*_ −*N*_*s*_), min(*K*_*s*_, *n*_*s*_) ⟧ . To test the association between features and outcomes, two cases are taken into account : the case where *k*_*s*_ are observed less than expected (*E*[*X*_*s*_]) (or more than expected), it is an under-association (or over-association) of positive outcomes with the feature.

Therefore, two tests can be constructed, respectively,

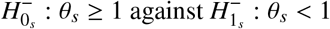

and

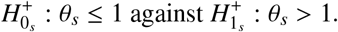

Where the *p*-value of the test is call respectively 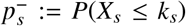 and 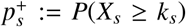, that are respectively the left and right tail of the distribution *X*_*s*_. Non-strict inequality is important to account for all possible tables up to the one observed. It is worth noting that 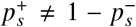, not being under-associated does not necessarily imply being over-associated. Finally, the one-tailed Fisher’s exact test (Fisher et al., 1936) has just been recalled here, also known as the hypergeometric test. The exact probability of observing one single table has been described. Now, the way to compute the exact probability of observing a set of table will be seen.

### 2.3. Stratified Truncated Fisher exact test

To assess the overall association between features and outcomes across different strata, since Fisher’s exact test describes the exact probability of observing one table, we can also describe the exact probability of observing a set of tables. In this approach, the *p*-values of each stratum are combined assuming independence between the strata, achieved by multiplying the *p*-values. The product may not be very pleasant to handle, which is why it is preferable to use its monotonic logarithmic transformation.

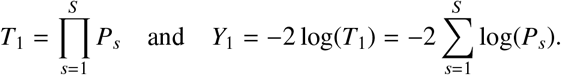

Where *P*_*s*_ is the random variable associated with the *p*-value of each independent test to be combined. Typically, this method is used to combine the *p*-values of tests with continuous distributions. In this case, so when the *p*-values follow a uniform distribution, the combination is the classic case of the combination by Fisher’s method (Fisher et al., 1936), where the combination *Y*_1_ follows a chi-squared distribution χ^2^ with 2*S* degrees of freedom.

The effect size can vary (e.g., efficiency of a drug that depends on the class of age of the patients) across strata and can even be in opposite directions (e.g., side effects taking over curation for a drug). Zaykin et al. (2002) and Zhang et al. (2020) introduced the idea of *p*-value truncation: by setting a cap and renormalizing *p*-values with respect to a user-defined significance threshold *τ*, the power of the test remains high and even avoids compensatory effects. The truncation threshold *τ* is a hyperparameter of the combination statistic that selects the significance threshold above which a stratum is taken into account in the statistic. The weighted truncated combination is given by :

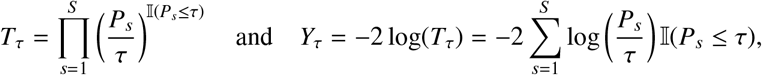

called soft-thresholding, where 𝕀 (·) is the indicator function. Always in the case where the *p*- values are uniformly distributed, Zhang et al. (2020) gives the expression of the survival function of *Y*_*τ*_ for any *τ* ∈ [0, 1].

When the *p*-values *P*_*s*_ come from Fisher’s exact test of each sub-table, they do not follow a uniform distribution, but a discrete distribution resulting from the hypergeometric test, which may be quite different from the distributions in the continuous case, also call sub-uniform distribution. The core of this paper presents a method for combining sub-uniform *p*-values.

Given that the combination *Y*_*τ*_ has a discrete distribution, the finite set of values taken by *Y*_*τ*_ can be explored to calculate the combined *p*-value exactly using an analytic solution. For a stratum *s*, Ω_*s*_ = ⟦max(0, *n*_*s*_ + *K*_*s*_ − *N*_*s*_), min(*K*_*s*_, *n*_*s*_) ⟧ is the support of the hypergeometric distribution, let us denote by the cumulative distribution function *F*_*X*_*s* (*x*) = *P*(*X*_*s*_ ≤ *x*) and the inclusive survival function 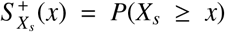, where *X*_*s*_ ∼ *HG*(*N*_*s*_, *n*_*s*_, *k*_*s*_). We define the Cartesian product Ω = Ω_1_ ×Ω_2_ ×· · · ×Ω_*S*_ of all possible realizatio ns o)f the tuple (*X*_1_, *X*_2_, …, *X*_*S*_), the support of the discrete random variable 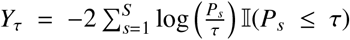 is the result of all possible value of Ω through the definition of *Y*_*τ*_. In our case, the cardinal of 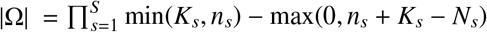

As before, there are two cases for the two one-sided *p*-values of each test. To study the overall under-association (or over-association) of positive outcomes with the feature, the *p*-values of each under-association (or over-association) test are combined. Combining the *p*-values involves studying the intersection of all the null sub-hypotheses.

For the overall under-association, the hypotheses are as follows :

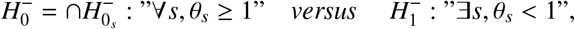

with the *p*-value, 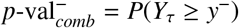 for the observed combination

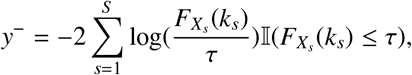

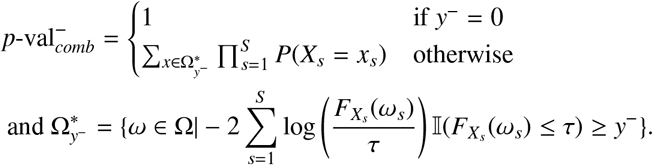

Where 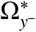 is the set of possible realizations in which the combination is greater than the observed combination *y*^−^.

Similarly, for the overall over-association :

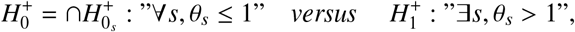

with the *p*-value, 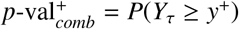 for the observed combination

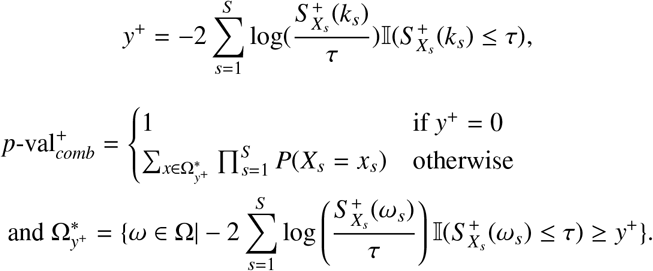

In summary, the combined *p*-value is calculated as the sum, until the observed combined value, of the product of the probability of the possible combinations of observations over all strata, even in the case of truncation where strata are not taken into account in the value of the combination test statistic. The *p*-values above the threshold are also taken into account when calculating the combined distribution. The considerations of numerical problems and cases of the same combination value for different realizations are provided in Supplementary Material B and Supplementary Figure 5.

As mentioned by Jung (2014); Chiba (2017), determining the exact distribution of *Y*_*τ*_ is computationally heavy. In fact, the set of all possible values of *Y*_*τ*_ must be explored to account for only realizations greater than the observed one. The size of this set is equal to the product of the size of the supports of each hypergeometric distribution. The complexity of the exact calculation therefore increases exponentially with the number of strata 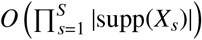. In some applications, the order of magnitude is 10^6^ 10^7^, and it is perfectly possible to explore all the possible combinations of the different strata and therefore obtain an exact calculation of the overall significance across strata. Our first contribution is a tool to compute this exact calculation. However, in other contexts, while maintaining the discrete nature of the *p*-values to be combined, the set of possible values may be excessively large, and it is impossible to calculate an exact value. In this case, rather than using an asymptotic test, which is a family of tests widely used in the literature, it is preferable to try to approximate the statistic of the exact test.

### 2.4. Gamma approximation

Given the potentially high computational cost of exact computation for the *p*-value of the stratified Fisher’s combination in discrete cases, we propose an approximation of the exact distribution followed by the statistic *Y*_*τ*_. As mentioned in the previous section, the approximation of *Y*_1_ by 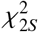, as in the case of uniformly distributed *p*-values on [0,1], is well known as Fisher’s method. However, through simulation, we observe that the distribution resulting from a combination of *p*-values from Fisher’s exact test differs slightly from a χ^2^ distribution in terms of scale and amplitude. Since the Gamma distribution Γ is a generalization of the χ^2^ distribution 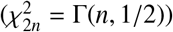, we approximate the discrete combination by finding *α* and *β* that make Γ(*α, β*) fit the distribution of *Y*_*τ*_. Through simulations, in Supplementary Material F with Supplementary Figure 10, we show that the gamma approximation asymptotically converges to the chi-squared distribution when the sample size *N*_*s*_ increases and random margins (∼ *U* ⟦1, *N*_*s*_⟧). This type of method has already been used in the literature to combine continuous dependent *p*-values (Brown, 1975; Kost and McDermott, 2002; Li et al., 2014; Poole et al., 2016; Yang et al., 2016; Zhang and Wu, 2023), and it has recently been used for Fisher’s combination of independent sub-uniform *p*-values (Contador and Wu, 2023). Indeed, Contador and Wu showed that the gamma distribution closest, in terms of Wasserstein distance, to any positive discrete random variable is the one that matches its mean and variance. They applied it to the Fisher’s combination of independent discrete *p*-values. Without truncation, the approximation is, therefore, as follows:

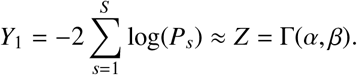

For the case where τ < 1, due to our truncation, the approximation of the truncated law of combination is in fact a mixture. The truncated combination law now contains a whole set of zero values corresponding to the case where all the combined *p*-values are greater than *τ* associated with *p*-val_*comb*_ = 1, which can be modeled by a Dirac distribution *δ* (whose survival function is a threshold function). And the rest of the distribution, in the case where at least one of the *p*-values is less than *τ*, is modeled by a Gamma distribution as before. The approximation model using a mixture distribution *Z* is therefore as follows :

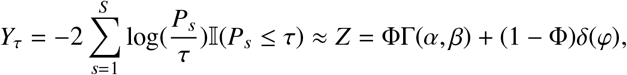

with 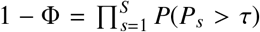 corresponding to the probability that all the *p*-values are greater than *τ* and the associated combination value.

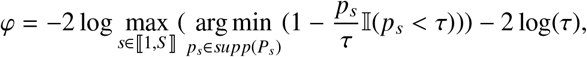

which corresponds to − 2 log of the largest *p*-value below the threshold *τ*.

The mixture involves a shift of *φ* in the survival function of Γ, writing *W* as the r.v. of Γ(*α, β*), the expression of the approximate combined *p*-value is given by :

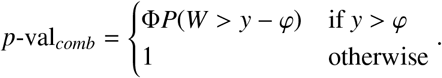

Inference of *α* and *β* is done by matching the moments of *Y*_*τ*_ and *Z* by analytically calculating the moments of the exact law of combination. Even though Cantador showed that the first and second moments are optimal in terms of the Wasserstein distance, we experimentally show in section 3.3 and in Supplementary Material A that matching higher orders moments may better control the type I error rate in some situations. The details of the computation of analytic moment matching using cumulants are also given in Supplementary Material A. The complexity of the gamma approximation is much lower than the exact calculation, since the exact moments of each stratum statistic *log*(*P*_*s*_) can be calculated analytically and combined linearly to obtain the moments of the overall statistic *Y*_*τ*_. The complexity is therefore linear in the number of strata 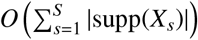.

### 2.5. Comparison to other statistical tests

Other statistical tests for assessing the overall significance between outcomes and features in stratified 2 × 2 contingency tables are documented in the literature. These tests vary in sensitivity and the conditions under which they are applied. We will compare our GASTE-test method with the following popular alternatives:

- Bonferroni’s procedure (Bonferroni, 1936) controls for the combined null hypothesis 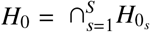 . At a threshold *α, H*_0_ is rejected if min 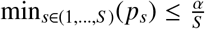. Although this procedure tends to be conservative, it exhibits power in the presence of a small number of strata (up to a single one) with strong effect.
- A widely used test in meta-analyses and often considered a reference method for stratified 2 × 2 contingency tables is the Cohran-Mantel-Haenszel test (CMH test) (Cochran, 1954; Mantel and Haenszel, 1959). Intermediate tests are not performed for each stratum; instead, data from all tables are used to construct a single test statistic based on the hypergeometric distribution. The test statistic is given by:

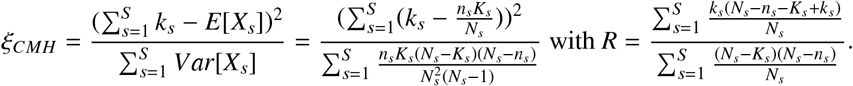 Testing whether the common odds ratio *R* differs from 1. Although this statistic rigorously follows a discrete distribution based on combinations of hypergeometric distributions, it is considered to asymptotically follow a χ^2^ distribution with 1 degree of freedom. However, the CMH test assumes homogeneity of the effect of the association between features and outcomes across strata. Opposite effects in different strata may cancel out, potentially escaping detection, due to the possible averaging effect in the sum of the common odds ratio. Therefore, the CMH test must be accompanied by the Breslow-Day test (Breslow et al., 1980) to verify this homogeneity assumption. Given its widespread use, comparing this method with our own, which does not rely on this homogeneity assumption, could be insightful.
- Classical Fisher’s *p*-value combination (Fisher et al., 1936) assumes uniformly distributed continuous *p*-values for each stratum. It has long been known that the Fisher’s combination is therefore inappropriate and can lead to false results if applied in discrete cases (Kincaid, 1962; Mielke Jr et al., 2004). Applied to hypergeometric tests, one thus ignores the discrete nature of the *p*-values. The 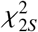 distribution followed by the combined statistic *Y*_*τ*_, in case of uniform *p*-values, will be compared with the exact and approximate distribution by GASTE to assess its deviation.

## 3. Results

### 3.1. Importance of exact calculation

In this section, we compare, under different scenarios, the exact distribution (analytically computed) of the statistic *Y*_*τ*_ with both the gamma approximation developed in this paper and the chi-square distribution used in Fisher’s combination method. To this end, five strata are simulated, resulting in a total number of combinations across five different tables of the order of magnitude 10^7^, with an average support size of 30 for each table. Specifically, we focus on cases where positive outcomes equal the number of features (equal marginals), we vary the number of samples *N*_*s*_ across four different contexts, so that the support size represents 50%, 20%, 10%, and 5% of the total, i.e., *n*_*s*_ = *K*_*s*_ = 0.5*N*_*s*_ for the first case. This deliberate selection places us in asymmetric scenarios in which study factors (positive outcomes and features) are not rare in the first two cases and can be considered rare in the latter two. Subsequently, for each of these four cases, five tables are simulated with an underlying hypergeometric distribution to conform to the null hypothesis context. Initially, we examine the non-truncated case (*τ* = 1) for this stratified table, evaluating the exact distribution (*Y*_1_), the uniform *p*-value approximation (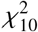), and the gamma distribution (Γ(*α, β*)) as a suitable approximation. As depicted in Figure 1, due to the asymptotic nature of the chi-square distribution, the rarer the features relative to the number of samples, the greater the difference between the exact value and the uniform *p*-value approximation (chi-squared). The Fisher chi-square combination appears conservative, while the gamma approximation fits the exact distribution very well in all cases. Indeed, the order of magnitude of the approximation error of the gamma distribution is a few percentage points, while the error of the chi-square distribution is on the order of a few hundred percentage points.

**Figure 1.**
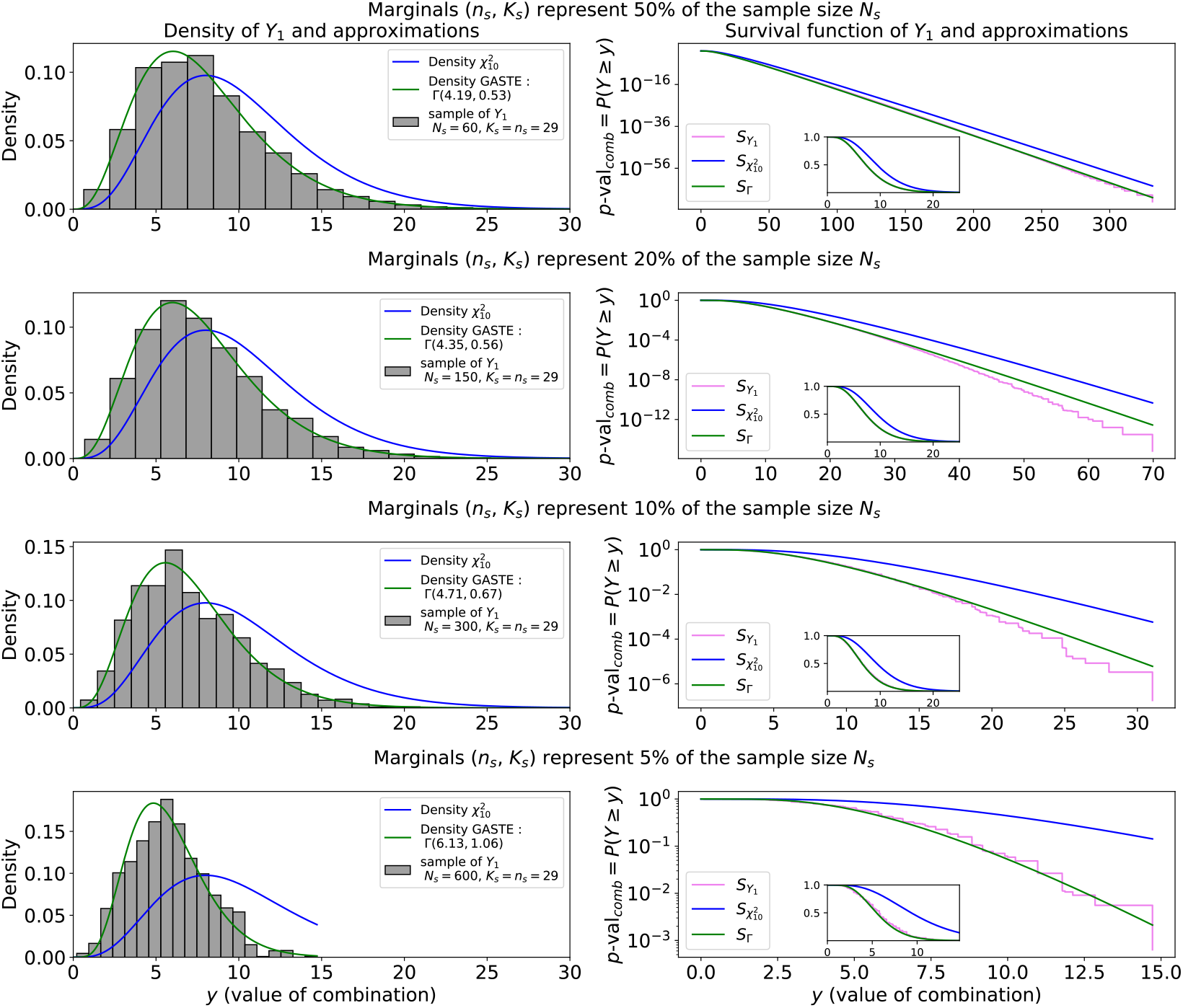
Comparison of stratified exact test, *Y*_1_, its gamma approximation, Γ, and χ^2^ distribution for the *p*-value combination of under-association. Study of four cases, depending on if features are rare or not, in the first case one in two samples has the features *N*_*s*_ = 60, *K*_*s*_ = *n*_*s*_ = 29, second case one in five *N*_*s*_ = 150, *K*_*s*_ = *n*_*s*_ = 29, third case one in ten *N*_*s*_ = 300, *K*_*s*_ = *n*_*s*_ = 29 and fourth case one in twenty *N*_*s*_ = 600, *K*_*s*_ = *n*_*s*_ = 29. On the right side the density function is plotted, the histogram is built from a sample of *Y*_1_ given by the exact calculation. And on the left side, the survival function of the distributions, i.e., the combined *p*-values for each possible *Y*_1_-combination value.

In this example, the number of possible combinations is exactly 24.3 millions, and the calculation time for the gamma approximation is 0.01*s* on a 13th Gen Intel® Core™ i7-13700H CPU compared to 55 seconds in parallel calculation on 20 CPUs of the same type. For a general comparison of the computation time, see Supplementary Figure 7 of the computation times with respect to the number of strata given in Supplementary Material D. For this simulation, the gamma approximation boosts the computation speed by more than a factor of 1000, while keeping the computation accurate (e.g., error *<* 6% against error *>* 300% for χ^2^, at a significant level of 10^−3^ for the first two cases). More details on the accuracy and computation time for the gamma approximation and χ^2^ are given in Supplementary Material C and Supplementary Figure 6.

The gamma approximation emerges as a suitable balance between precision and computational efficiency. This shows the importance of performing exact tests in the case of sub-uniform *p*-values and that the gamma approximation is a good general approximation.

### 3.2. Power of the test on simulations

Having shown that approximating the stratified exact test is essential to reduce computation time considerably, we can now run simulations to compare it with other tests. A second study was therefore carried out to compare the power of different tests and their behavior in homogeneous or non-homogeneous cases. We will compare our gamma approximation method of the stratified truncated exact test (GASTE) for three truncations *τ* = 1 (no truncation), *τ* = 0.2, *τ* = 0.05 with the Bonferroni test, the CMH test and the classic Fisher’s combination following a χ^2^ degree of freedom 2*S* .

For the simulation parameters, to avoid having identical marginals in each stratum and create small variation in the input data, we draw the marginals according to a Poisson distribution with parameter *µ* = 30 and then consider them as fixed and to remain in the case of our conditional tests. Four strata are arbitrarily chosen, each with a sample size of *N*_*s*_ = 150, to be in the conditions of the second scenario described in the first simulation (previous section). This scenario involves a relatively minor rare factor compared to the sample size (≈ 20%), which could reflect a realistic data case, and the choice is not biased toward favoring the GASTE method over the other method, as we do not opt for conditions where the method performs best (e.g., low marginals and large sample size).

We create 500 different sets of 4 tuples (*N*_*s*_, *n*_*s*_, *K*_*s*_) to define the sample size and the marginals of 500 stratified contingency tables of 4 strata. For these 500 sets of stratified table parameters, we apply an association effect to each stratum to reject the null hypotheses 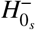 and 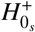 at a given significance level. When simulating positive (or negative) effects, the probability, under the null hypothesis, of drawing an individual belonging to the class of positive outcomes is multiplied (or divided) by the chosen effect size. Based on this new probability, the number of positive outcomes with features (*k*_*s*_) is drawn. This generates an over-association (or under-association) of positive outcomes with the feature. Subsequently, various global significance tests are applied to compare their power. Where the effect size is positive (or negative), the global over-association (or under-association) is studied. For a chosen effect size, and for each of the 500 simulated marginal tuples, we simulate 50 contingency tables (such that only *k*_*s*_ is redrawn). For one effect size, we therefore run 25,000 stratified table simulations (50× 500) and as many global significance tests. Figure 2 illustrates the results of this study.

**Figure 2.**
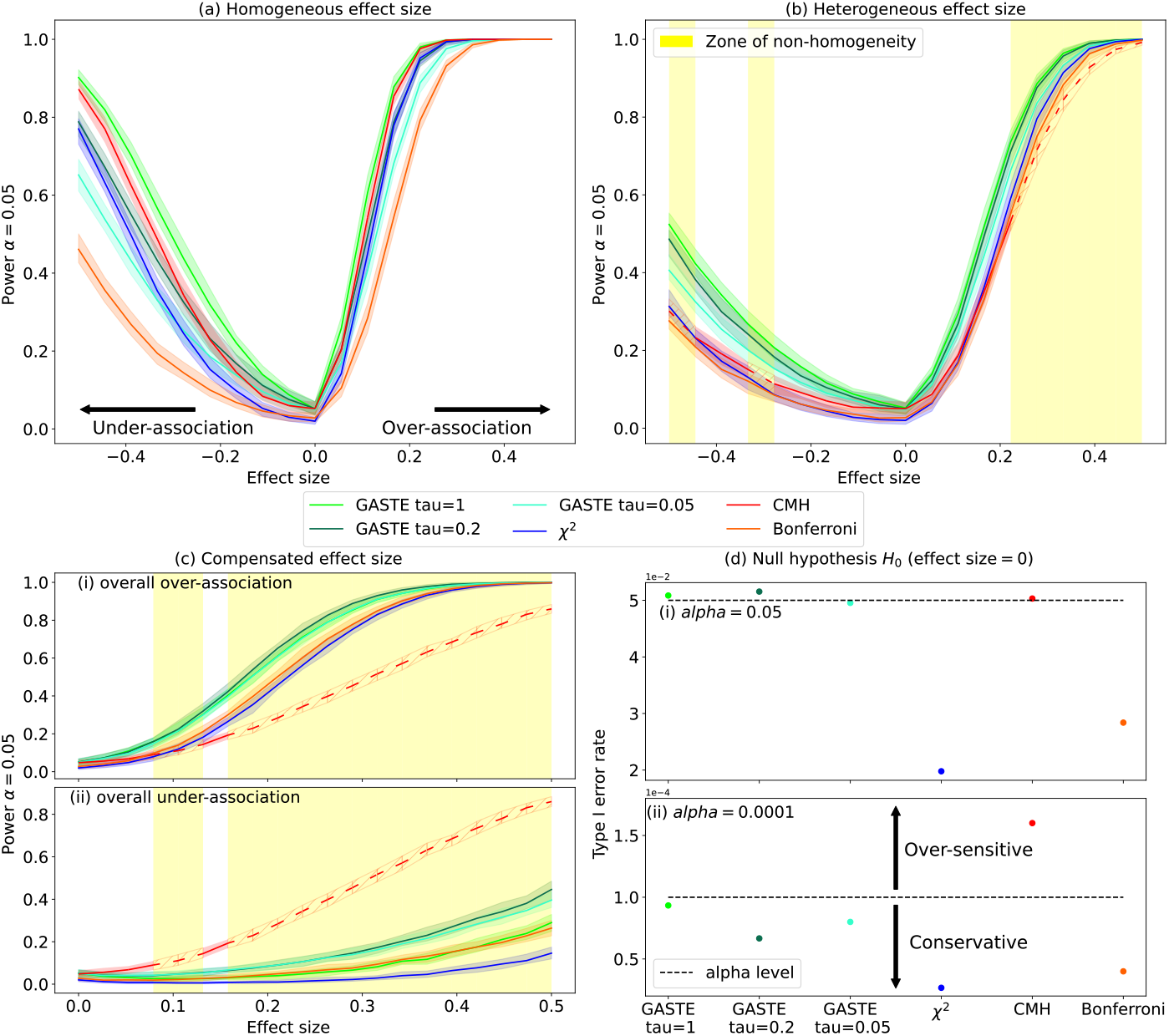
Study of 3 different effect size configurations, the first configuration, sub-figure (a), corresponds to a homogeneous case where the effect size is applied to the all 4 strata in the same way. The second configuration, sub-figure (b), corresponds to a case of heterogeneity where the effect size is only applied to two strata and the other two are in null condition. The third configuration, sub-figure (c), corresponds to a case of compensation where the effect size is applied positively to two strata and negatively to the other. For these configurations, the areas where the significance of the strata is not homogeneous using the Breslow-Day test are illustrated by a yellow background. In this zone, the CMH test appears as a hash because it should only be applied if the homogeneity test is satisfied. In sub-figure (d), the type I error is studied for significant levels alpha, 0.05 and 10^−4^ such that under null hypothesis (effect size of all strata are null) the three configurations are concatenated to obtain 75,000 simulations

In cases of homogeneity, our gamma approximation method without truncation demonstrates comparable, or slightly superior, power relative to the CMH test (16% more powerful on average). Unsurprisingly, the Bonferroni test is the most conservative and the χ^2^ test is less powerful than GASTE. In instances of heterogeneity, where the CMH test cannot be applied due to strata heterogeneity for some effect sizes, our GASTE method emerges as the most powerful (54% more powerful on average on effect size outside non-homogeneity). Given that the marginals *n*_*s*_ and *K*_*s*_ are less than half the sample size *N*_*s*_, for the same absolute effect size, we observe weaker statistical power for under-association compared to over-association, owing to the natural rarity of positive outcomes with the feature. GASTE proves particularly potent in scenarios involving multiple small effects spread across strata. In instances of compensation, truncated GASTE methods exhibit greater power compared to the non-truncated approach. This is because in cases of compensation, certain strata exhibit significant effects in opposite directions, which truncation helps to mitigate by removing opposing effects while retaining significant ones. Furthermore, the use of the CMH test in compensation scenarios is erroneous; in our case, due to the asymmetric margins relative to the sample size, the effects of compensation are not balanced. This falsely inflates the apparent importance of under-association because over-association is inherently easier to detect. This underscores the importance of separately studying over-association and under-association in non-homogeneous strata, as well as the necessity of verifying assumptions before applying the CMH test to avoid misleading conclusions. Finally, in the case of null effect sizes, all tests remain below the significant level *α* = 0.05. However, for a significant level *α* = 10^−4^, the CMH test demonstrates over-sensitivity, leading to inflated significance, even under null conditions. This highlights an additional concern with the CMH test, as lower significant levels correspond to higher false positive rates, posing challenges for its reliability, particularly if the *p*-value of the CMH test is used in a multiple testing context.

### 3.3. Choice of moments through type I error control study

In the gamma approximation, we infer the parameters *α* and *β* by analytically matching the moments of the exact distribution of the combination *Y*_*τ*_, and the moments of the gamma distribution *Z*. We can naturally match the mean and variance, but despite Contador and Wu (2023) work showing that matching the first two moments minimize the Wasserstein distance between the continuous gamma distribution and the discrete combination distribution, in some cases it may be interesting to choose a higher order moment to better control the type I error rate. Indeed, presented in Supplementary Material A with Supplementary Figures 2-4 where we perform large simulations as a function of truncation *τ*, sample size *N*_*s*_, marginals *n*_*s*_ and *K*_*s*_, and the alpha level of type I error control, matching the variance or excess kurtosis (fourth order moment) leads to better control of the type I error rate at certain alpha levels. In the case of a statistical test, it is more interesting to control the lower quantiles than to have the best control over the whole distribution on average. Criteria based on the Wasserstein distance may not be the most relevant in this type of application. In general, using moments of order one and two is a good choice that we recommend by default; however, for high levels of significance, i.e., very low alpha levels, if the tail distribution of the combination law is very spread out, it may be interesting to choose a higher-order moment that does not minimize the distance between the distributions on the whole distribution, but more on the tail distribution. The level of truncation also influences this point, so that the higher-order moment is better suited to high levels of truncation. We found that an empirically good indicator of the spread of the tail distribution is the ratio between the variance of the combination law and the largest possible value taken, i.e., *Var*[*Y*_*τ*_]*/* max(*supp*(*Y*_*τ*_)). If this ratio is low, the tail of the distribution is very spread out. This also corresponds to cases where the maximum achievable level of significance is very low (smallest possible combined *p*-value).

We suggest the following strategy for using GASTE: by default, we propose to calculate the exact combination test if it can be calculated in a suitable time (typically *supp*(*Y*_*τ*_) =| Ω| < 5.10^7^, adjustable parameter). Otherwise, the gamma approximation is proposed with a truncation threshold *τ* = 0.2 which seems reasonable, for example, to set aside strata without an association signal. With this default value of *τ*, the default fourth-order moment is chosen for the gamma approximation. But the user is free to set any value of *τ* and any second, third, or fourth order moment. If the user sets a *τ* greater than 0.2, or if the tail of the distribution is not spread out (^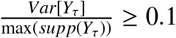^), we automatically switch to using a moment of order two.

### 3.4. Application to real datasets

#### 3.4.1. Application I, Detection of associations between plant species

We begin by describing the application that motivated the development of our method, using the ORCHAMP pinpoints dataset (Thuiller et al., 2024). Our dataset comprises information on the presence or absence of plants (1,048 different species) across various observation zones (177 zones) in the French Alps, each considered independent. The goal is to ascertain whether the observed coexistence between two species differs significantly from what would be expected under a null model, knowing that they are seen in the same observation zone (co-occurring in the same environment). To achieve this, 2 × 2 contingency tables are used, where rows denote the presence or absence of one species and columns denote the presence or absence of the other. One-sided Fisher’s exact test is used to evaluate the independence between the two species in a given environment. Subsequently, our GASTE method is employed to derive global insights across all environments, with each environment treated as an independent stratum.

Each observation zone consists of approximately 300 survey points in a predefined 30 meters × 30 meters area with the same survey protocol, with some species being highly prevalent (present more than 250 times out of 300) while others are rare (present only a few times). For every pair of plants in the dataset, the exact combined *p*-value is computed if the support of *Y*_*τ*_ is less than five million combinations; otherwise, a gamma approximation with the fourth-order moment is used. To reduce the number of pairs of plants to study, a filter is applied to the data by excluding pairs where the smallest achievable exact level of significance is over 0.2. Indeed, for example, it is not feasible to study the under-association of two rare species, as the probability of not observing any point with both plants is already high by nature. In total, 63749 pairs of plants are studied in over-association and 10513 in under-association. Multiple testing corrections are applied when examining all plant pairs of interest, with a Bonferroni correction utilized to identify significant positive and negative plant pair associations. Moreover, a truncation threshold of *τ* = 0.2 is applied to increase the statistical power of the test, to cope with the potential heterogeneity of the association signal between plants among environments.

Similarly to the approach illustrated in Figure 2, we compare the detection of coexistence using the various tests mentioned above on real ORCHAMP data. This comparison is visualized through a Venn diagram (Figure 3) and under the null model (Table 3.4.1). The null model was obtained by randomly distributing the species 400 times over the survey points while maintaining the abundance of each species in each observation zone.

**Figure 3.**
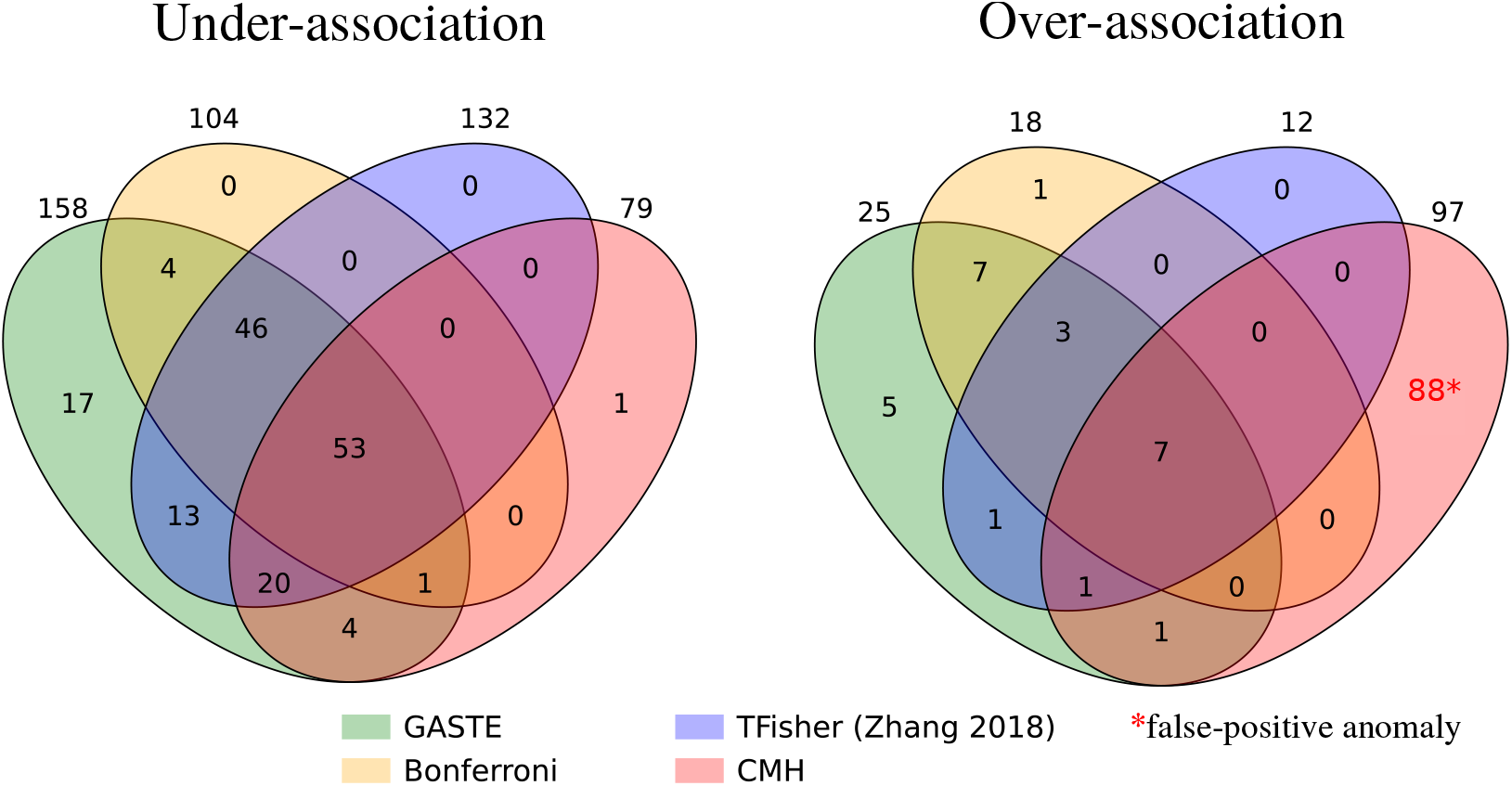
Venn diagram of the detection of association between two plants species in the ORCHAMP data. The first diagram represents the detection of under-association, the second diagram represents the detection of over-association. The different tests are represented by the different circles, the Bonferroni test, the CMH test, the TFisher test and our GASTE method. Note that the over-associations detected by the CMH method are mostly spurious detections due to rare species counts, as witnessed by our FWER assessment.

Figure 3 illustrates that the GASTE method detects the most pairs (≈ 20% more than TFisher (uniformity assumption) and twice as many as CMH) while maintaining the probability of making at least one type I error (FWER) under the significant level *α* (Table 3.4.1). More precisely, our GASTE method detects 158 pairs of plants significantly under-associated and 25 pairs significantly over-associated across the different zone of observation, means co-occurring in the same zone (environment) but respectively significantly not coexisting and significantly coexisting in the same survey point. These two sets overlap in 6 plant pairs, meaning that the two plants are significantly under- and over-associated at the same time. This does not lead to an erroneous result, but shows that the association between the two plants depends on the environment, as a harsh environment leads plants to evolve in the same restricted area, and inversely, a rich environment leads plants to expand and conquer an area, producing competition between plants. A more precise example is given in Supplementary Material E.1 with Web Figure 8 on the association between *Agrostis schraderiana* and *Rumex arifolius* species, concluding that the pair brings a significantly overall signal of under-association in an open environment such as grasslands and brings a significantly overall signal of over-association in a closed environment such as forest.

An unusual observation in the null model results is the family-wise error rate of the CMH test. Firstly, this anomaly is asymmetrical; it occurs in the case of over-association but not under-association. Secondly, the error rate is significantly higher than the significant level *α* for all levels tested, as it is always equal to 1. This means that during each randomization, more than one pair of plants is detected in over-association under the null hypothesis. These detections correspond to cases where the marginals are very small, just a few counts, which explains why they are found only in over-association. Our pre-selection of pairs prevents them from being studied in under-association, but it is possible to evaluate the over-association of two rare species. However, in the case of the CMH test, these pairs do not comply with Cochran’s rule, which states that the counts in the contingency tables must each be greater than 5. Consequently, the detection of over-association by the CMH method exceeds that of the other methods, with 88 pairs detected. These detections are very likely false positives that do not adhere to the Cochran rule, and this hypothesis would need to be verified for each pair. This highlights the problem of the over-sensitivity and generality of the CMH test, particularly in cases where the marginals are low.

In addition, excluding this anomaly, the FWERs are relatively lower than the *α* significance level due to the Bonferroni correction in multiple tests. However, once again, our GASTE method has a better detection rate than the other methods, while the conservative nature of the Bonferroni and TFisher methods is salient.

#### 3.4.2. Application II, Berkeley’s admission (1973)

For this second application, we take a widely used example of stratified data that illustrates Simpson’s paradox (Simpson, 1951) and highlights the value of stratifying data. The dataset known as UCBAdmissions lists admissions to the University of California, Berkeley, in 1973 according to gender and by department (from A to F). The study of these data was popularized by the apparent discrimination against women in admissions (Bickel et al., 1975). In fact, comparing the overall admission rate, 44% of men were admitted (1198 out of 2691) compared to 30% of women (557 out of 1835). However, by stratifying the data by department, this trend tends to be reversed.

For each department, a 2× 2 contingency table is created, with men and women in the columns and admissions and refusals in the rows. The association studied is therefore men and admissions, in order to detect whether there are more or fewer men admitted than in the case of a null model (independence between sex and admission). We recreated a Forest plot (Figure 4) summarizing the data from the tables stratified by department (odds ratio, confidence interval, CMH weights), and added the information from our two global over and under association tests, as well as the popular CMH test and the Breslow-Day homogeneity test. The homogeneity criterion is not fulfilled (Breslow-Day test *p*-value 0.002) and prevents the use of the CMH test. However, as was done in the original study without checking homogeneity, the overall association with CMH test is announced as non-significant (*p*-value 0.22). The non-homogeneity of the strata already distorts the result of the CMH test. Moreover, in department A there is a strong under-association of admissions by men compared with women (*p*-value Fisher’s exact test 1.15e-5). This under-association in this stratum (department) drives the overall under-association test, making it also significant (*p*-value_*combined*_ 0.0012). The result of our GASTE test is therefore that if men and women had been admitted to the departments in the same way, the probability of observing such results is about one in a thousand. By artificially adding departments with results similar to departments B to F, for it to be likely to observe such a difference in department A at random (*p*-value_*combined*_ *>* 0.05), there would have to be 20 departments in total. These results were obtained using our GASTE approximation method, as the exact calculation would require evaluating trillions of combinations.

**Figure 4.**
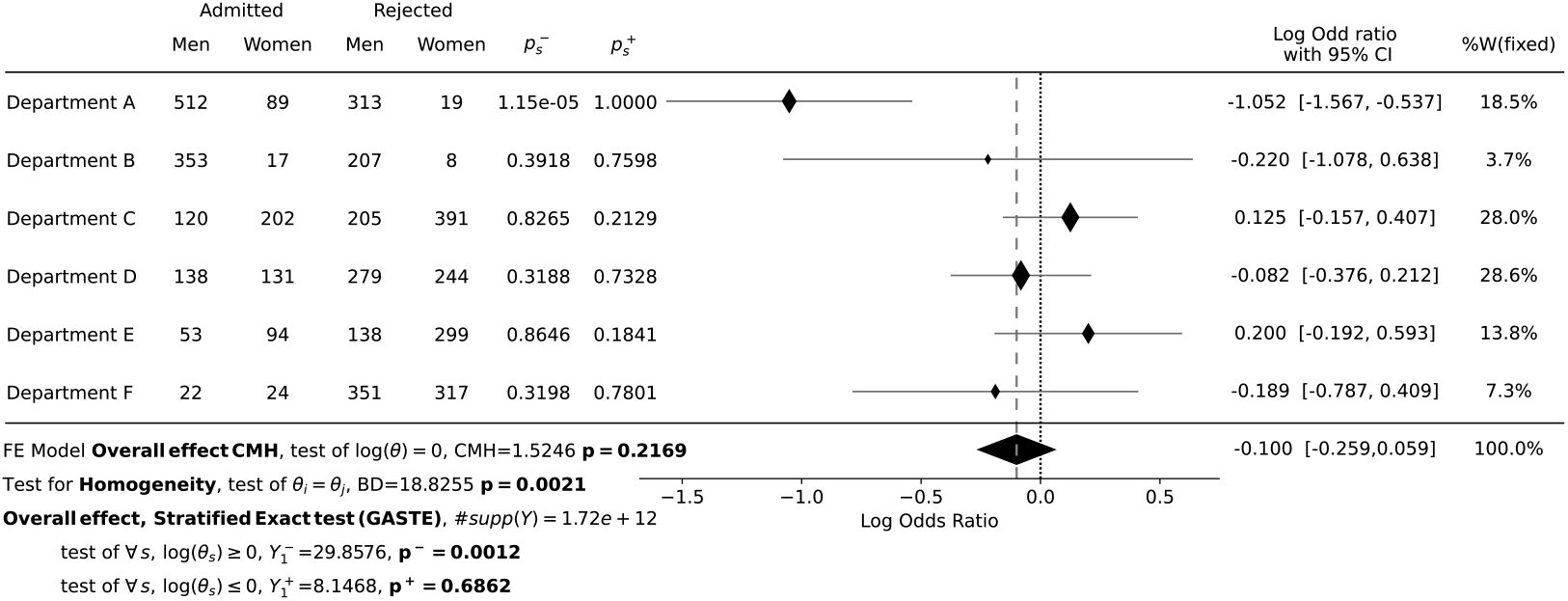
Berkeley’s admission (1973) stratified analyse

In conclusion, while there appears to be an overall under-association of men compared to women in admissions, this does not necessarily indicate a gender bias in admissions. It is possible that women outperformed men in department A to such an extent that it created the perception of an overall under-association of men.

## Discussion

We believe that our GASTE method offers substantial value in the statistical analysis of stratified binary data. As demonstrated in both our simulations and on real data applications, the GASTE method is a powerful test that operates without assuming homogeneous associations between features and outcomes across strata and without a minimum data requirement in the contingency tables. This contrasts with the CMH test, which, despite its widespread use, suffers from both of these limitations. Applying statistical tests such as CMH test without meeting their assumptions (e.g., homogeneity, uniformity of the *p*-values for Fisher’s combination, sufficient counts for CMH) can lead to both false discoveries and undetected effects. Drawing general rules about the approximate fulfillment of underlying assumptions is difficult in practice: as shown in the simulation (cf. Figure 1, bottom plot), when the marginals are low, the asymptotic test is far from being accurate even with large sample sizes (e.g., even with *N >* 600). To avoid these limitations, it is crucial to conduct exact tests whenever possible, or use approximate computations of the exact statistic when computationally intractable, offering high accuracy at low computational cost (Supplementary Material C and Supplementary Figure 6). The GASTE method performs effectively in scenarios with both homogeneous and heterogeneous effects between strata, and in cases of both low and high significance. The major point of our method is to approximate the exact test rather than relying on asymptotic tests, enabling a wide variety of applications thanks to the computational efficiency. Our method can trigger detection as soon as odds ratio of a single stratum is sufficiently far from 1 (i.e., the stratum has a sufficiently significant *p*-value) compared with the other strata, as well as when the signal is diffuse (i.e., in each stratum the *p*-value has a low level of significance, but this low significance is found across several strata), for example Application I, Supplementary Material E.2, Supplementary Figure 9. Computing the exact probability of observing a set of 2× 2 contingency tables under the null model can be straightforwardly rephrased into a problem of combining the discrete *p*-values of each stratum. This formulation, in turn, led us to naturally propose an approximation of the exact test. Unlike usual methods, we believe it is essential to consider under-association and over-association separately (rather than using two-sided *p*-values), as in cases of heterogeneity (cf Application I and Supplementary Material E), both associations can be significant simultaneously, providing a more refined analysis. Incorporating truncation into the *p*-value combination enhances statistical power in scenarios featuring few effects or contradictory effects between strata. However, when effects are homogeneous, truncation may not augment power and could potentially diminish it. Thus, truncation should be considered beforehand depending on the considered problem. If the nature of the signal to be detected is completely unknown, instead of choosing an arbitrary *τ* value, an omnibus test could be constructed from several GASTE tests with different values of *τ*. This approach smooths the detection power across truncations. As these *p*-values are obviously dependent, an interesting avenue of research would be to check the suitability of the uniformly weighted Cauchy combination (Liu and Xie, 2020) for the case of sub-uniform *p*-values.

Moreover, we think our method will be of great interest for many applications. In the case of meta-analysis using 2 × 2 tables, each stratum corresponding to a distinct and independent study, thus verifying the hypothesis of independence between strata, our method does not require selecting a set of studies that comply with homogeneity criterion, thus avoiding the risk of “cherry picking” for meta-analysis (Yoneoka and Rieck, 2023). This mitigates both study selection bias (Felson, 1992; Eisend and Tarrahi, 2014), particularly in heterogeneous scenarios, and publication bias (Rothstein et al., 2005; Kicinski et al., 2015), since even studies with non-significant results provide information within a multiple-testing or *p*-value combination framework. We also posit that our method holds promise in epidemiology. Here, each stratum corresponds to a confounding clinical feature of independent patients in which sample sizes (i.e., marginals) are not always controllable and can turn out to be low. Even if our method performs well in general, there are still some pitfalls when using the gamma approximation. Despite the optimality guarantee on matching the first two moments for the Gamma distribution in terms of Wasserstein distance, there is no guarantee that it is optimal in terms of the tail of the distribution, and thereby for controlling the type I error rate effectively. That is why we implemented higher-order moments that empirically show improvements in type I error control for light-tailed situations (Section 3.3 and simulation Supplementary Material A). For further research, it would be interesting to find theoretical guarantees on the best moment to choose for a given problem. Finally, the GASTE method focuses on the combination of *p*-values stemming from the hypergeometric distribution, but the same idea both for computing the exact law and for the gamma approximation can easily be transferred to other discrete laws with finite support.

## Conclusions

We have reformulated the stratified exact test problem as a Fisher’s combination of one-sided sub-uniform *p*-values. We propose an exact algorithm as well as a gamma approximation of the exact combination statistic for cases in which exact calculation is prohibitively expensive. Furthermore, we have extended this test and its approximation to incorporate truncation, thereby enhancing statistical power in scenarios with non-homogeneous and opposing effects between strata. We provide some empirical advice on the choice of the parameters of the approximation. The GASTE method offers a general approach suitable for both lowand high-significance cases in homogeneous and heterogeneous settings. Compared to the conventional CMH test, our method appears to be more suitable for studies with stratified 2 × 2 tables.

To use the method in practice, we have developed a Python package (easily wrappable in R) available on PyPI at https://pypi.org/project/gaste-test/ (open-source code available at https://github.com/AlexandreWen/gaste).

## Supplementary Material

**Supplementary Material A : Method of moment estimator for the gamma approximation :** Presentation of calculation of analytical moments of the exact distribution of combination *Y*_*τ*_ for 2nd, 3rd and 4th moment and inference of *α* and *β* parameters of gamma distribution. Results of the type I error control study for different moments, truncation *τ* values and parameters of contingency tables.

**Supplementary Material B : Numerical issue of exact calculation of the combined** *p***value :** Consideration of limit representation of floating point number in the exact calculation of the combined *p*-value and cases of same combination value for different realizations.

**Supplementary Material C : Time-accuracy simulation 1 :** Presentation of time of computation and accuracy of the gamma approximation and χ^2^ distribution presented in Figure 1.

**Supplementary Material D : Figure time complexity :** Computational time of the gamma approximation and exact calculation with respect to the number of strata.

**Supplementary Material E : Example of Forest plot on Orchamp data :** Presentation of the generality of GASTE through compensatory and widespread signals example in the ORCHAMP data.

**Supplementary Material F : Asymptotic convergence of Gamma model :** Inference of the *α* and *β* gamma parameters with respect to the sample size.

The data and the code to reproduce all results in the papers are available at https://gricad-gitlab.univ-grenoble-alpes.fr/wendlina/gaste_method.

