## Supplemental File for "Gamma Approximation of Stratified Truncated Exact test (GASTE-test) & Application"

### Supplementary Materials for Gamma Approximation of Stratified Truncated Exact test (GASTE-test) & Application

Univ. Grenoble Alpes, CNRS, Grenoble INP, LJK, 38000 Grenoble, France

#### Material A : Method of moment estimator for the gamma approximation

We recall the approximation model for the truncated combination law. For the case where $\tau < 1$, due to our truncation, the approximation of the truncated law of combination is in fact a mixture. The truncated combination law now contains a whole set of zero values corresponding to the case where all the $p$-values are greater than $\tau$ associated with $p\text{-val}_{comb} = 1$, which can be modelled by a Dirac distribution $\delta$ (survival function of the *Dirac* is a threshold function). And the rest of the distribution, in the case where at least one of the $p$-values is less than $\tau$, is modelled by a Gamma distribution as before. The approximation model by a mixture distribution $Z$ is therefore as follows:

$$Y_\tau = -2\sum_{s=1}^{S} \log\left(\frac{P_s}{\tau}\right) \mathbb{I}(P_s \leq \tau) \approx Z = \Phi\Gamma(\alpha, \beta) + (1-\Phi)\delta(\varphi)$$

with $1 - \Phi = \prod_{s=1}^{S} P(P_s > \tau)$ corresponding to the probability that all the $p$-values are greater than $\tau$ and the associated combination value $\varphi = -2\log \max_{s\in[\![1,S]\!]} ( \operatorname*{arg\,min}_{p_s \in supp(P_s)} (1 - \frac{p_s}{\tau}\mathbb{I}(p_s < \tau))) - 2\log(\tau)$ which corresponds to $-2\log$ of the largest $p$-value below the threshold $\tau$.

The parameters $\alpha$ and $\beta$ are inferred by matching the moments and here the mixture involves a shift of $\varphi$ in the survival function of $\Gamma$, so as if we write $W$ as the r.v. of $\Gamma(\alpha, \beta)$ the expression of the approximate combined $p$-value is given by :

$$p\text{-val}_{comb} = \begin{cases} \Phi P(W > y - \varphi) & \text{if } y > \varphi \\ 1 & \text{otherwise} \end{cases}$$

We begin by calculating the moments of order 1 and 2 of $Y_\tau$ (expectation and variance) and we will explain the calculation by detailing the random variables $P_s$ by the random variables $X_s \sim HG(N_s, n_s, k_s)$ by putting ourselves in the case of a test of under-association (the case of over-association is completely analogous by replacing $P(X_s \leq k)$ by $P(X_s \geq k)$).

##### 2nd moment

$$\begin{aligned}
\mu = E[Y_\tau] &= E[\sum_{s=1}^{S} -2\log\left(\frac{P_s}{\tau}\right)\mathbb{I}(P_s \leq \tau)] \\
&= -2\sum_{s=1}^{S} E[\log\left(\frac{P_s}{\tau}\right)\mathbb{I}(P_s \leq \tau)] \\
&= -2\sum_{s=1}^{S}\sum_{k\in supp(X_s)|P_s\leq\tau} P(X_s = k)\log\left(\frac{P(X_s \leq k)}{\tau}\right)
\end{aligned}$$

$$\begin{aligned}
\sigma^2 = Var[Y_\tau] &= Var[\sum_{s=1}^{S} -2\log\left(\frac{P_s}{\tau}\right)\mathbb{I}(P_s \leq \tau)] \\
&= 4\sum_{s=1}^{S} Var[\log\left(\frac{P_s}{\tau}\right)\mathbb{I}(P_s \leq \tau)] \qquad \text{by independence of } p\text{-value} \\
&= 4\sum_{s=1}^{S} E[\log\left(\frac{P_s}{\tau}\right)^2\mathbb{I}(P_s \leq \tau)] - E[\log\left(\frac{P_s}{\tau}\right)\mathbb{I}(P_s \leq \tau)]^2 \\
&= 4\sum_{s=1}^{S}\sum_{k\in supp(X_s)|P_s\leq\tau} P(X_s = k)\log\left(\frac{P(X_s \leq k)}{\tau}\right)^2 \\
&\quad - \left[\sum_{j\in supp(X_s)|P_s\leq\tau} P(X_s = j)\log\left(\frac{P(X_s \leq j)}{\tau}\right)\right]^2
\end{aligned}$$

We have everything we need to calculate the moments of $Y_\tau$, so let's look at those of $Z$:

$$E[Z] = \Phi E[Gamma(\alpha,\beta)] + (1-\Phi)\underbrace{E[\delta(\varphi)]}_{=0} = \Phi\alpha\beta$$

To calculate the variance, we use the formula for the total variance on the mixture:

$$\begin{aligned}
Var[Z] &= \Phi Var[Gamma(\alpha,\beta)] + (1-\Phi)\underbrace{Var[\delta(\varphi)]}_{=0} \\
&\quad + \Phi(1-\Phi)E[Gamma(\alpha,\beta)]^2 + \Phi(1-\Phi)\underbrace{E[\delta(\varphi)]^2}_{=0} \\
&= \Phi\alpha\beta^2 + \Phi(1-\Phi)\alpha^2\beta^2
\end{aligned}$$

By matching the moments we end up with the following system:

$$\begin{cases} E[Y_\tau] = E[Z] \\ Var[Y_\tau] = Var[Z] \end{cases}$$
$$\iff \begin{cases} \mu = \Phi\alpha\beta \\ \sigma^2 = \Phi\alpha\beta^2 + \Phi(1-\Phi)\alpha^2\beta^2 \end{cases}$$
$$\iff \boxed{\begin{cases} \alpha = \frac{\mu^2}{\Phi\sigma^2-(1-\Phi)\mu^2} \\ \beta = \frac{\Phi\sigma^2-(1-\Phi)\mu^2}{\Phi\mu} \end{cases}}$$

##### 3rd moment

We can also inferre the shape parameter $\alpha$ of the Gamma distribution using the third-order standardized moment, know as skewness ($Skew[Y_\tau]$). To simplify the notation, we now use the notation $Y_\tau = \sum_{s=1}^{S} W_s$ with $W_s = -2log\left(\frac{P_s}{\tau}\right)\mathbb{I}(P_s \leq \tau)$. By definition, the skewness of a random variable $Y_\tau$ is given by the following formula:

$$\gamma_1[Y_\tau] = Skew[Y_\tau] = E\left[\left(\frac{Y_\tau - E[Y_\tau]}{\sqrt{Var[Y_\tau]}}\right)^3\right] = \frac{E[(Y_\tau - E[Y_\tau])^3]}{Var[Y_\tau]^{3/2}} = \frac{\kappa_3[Y_\tau]}{Var[Y_\tau]^{3/2}}$$

where $\kappa_3[Y_\tau]$ is the third-order cumulant of $Y_\tau$. We use the property of cumulants, which is that the cumulant of a sum of independent variables is equal to the sum of the cumulants of each variable in the sum. So $\kappa_3[Y_\tau] = \sum_{s=0}^{n} \kappa_3[W_s]$. Finally, we have :

$$\gamma_1[Y_\tau] = \frac{\sum_{s=1}^{S}\left(E[{W_s}^3] - 3E[W_s]E[{W_s}^2] + 2E[W_s]^3\right)}{Var[Y_\tau]^{\frac{3}{2}}}$$

$$\text{where } E[{W_s}^n] = (-2)^n \sum_{k \in supp(X_s)|P_s \leq \tau} P(X_s = k)\log\left(\frac{P(X_s \leq k)}{\tau}\right)^n$$

We have described the skewness of $Y_\tau$, so now we want to calculate the skewness of $Z$ to match it. By showing that $E[Gamma(\alpha,\beta)^n] = \frac{\beta^n\Gamma(n+\alpha)}{\Gamma(\alpha)}$ and using the fact that all moments of a *Dirac* are zero, we show that $E[Z^n] = \Phi\frac{\beta^n\Gamma(n+\alpha)}{\Gamma(\alpha)}$, so we have :

$$\begin{aligned}
\gamma_1[Z] = E\left[\left(\frac{Z - E[Z]}{\sqrt{Var[Z]}}\right)^3\right] &= \frac{E[Z^3] - 3E[Z]E[Z^2] + 2E[Z]^3}{Var[Z]^{3/2}} \\
&= \frac{\Phi\frac{\beta^3\Gamma(3+\alpha)}{\Gamma(\alpha)} - 3\Phi\alpha\beta\frac{\beta^2\Gamma(2+\alpha)}{\Gamma(\alpha)} + 2(\Phi\alpha\beta)^3}{[\Phi\alpha\beta^2(1+\alpha(1-\Phi))]^{\frac{3}{2}}} \\
&= \frac{\Phi\beta^3(2+\alpha)(1+\alpha)\alpha - 3(\Phi\alpha\beta)^2\beta(1+\alpha) + 2(\Phi\alpha\beta)^3}{[\Phi\alpha\beta^2(1+\alpha(1-\Phi))]^{\frac{3}{2}}} \\
&= \frac{(2+\alpha)(1+\alpha) - 3\Phi\alpha(1+\alpha) + 2(\Phi\alpha)^2}{\sqrt{\Phi\alpha}[1+\alpha(1-\Phi)]^{\frac{3}{2}}}
\end{aligned}$$

$$\iff$$

$$\begin{aligned}
\gamma_1[Z]^2 &= \frac{\left[(2+\alpha)(1+\alpha) - 3\Phi\alpha(1+\alpha) + 2(\Phi\alpha)^2\right]^2}{\Phi\alpha[1+\alpha(1-\Phi)]^3} \\
&= \frac{4 + (12-12\Phi)\alpha + (13-30\Phi+17\Phi^2)\alpha^2 + (6-24\Phi+30\Phi^2-12\Phi^3)\alpha^3 + (1-6\Phi+13\Phi^2-12\Phi^3+4\Phi^4)\alpha^4}{\Phi\alpha + 3\Phi(1-\Phi)\alpha^2 + 3\Phi(1-\Phi)^2\alpha^3 + \Phi(1-\Phi)^3\alpha^4}
\end{aligned}$$

This gives us the following system:

$$
\begin{cases} E[Y_\tau] = E[Z] \\ \gamma_1[Y_\tau] = \gamma_1[Z] \end{cases}
$$

$$
\iff \boxed{\begin{cases} \beta = \frac{E[Y_\tau]}{\Phi\alpha} \\ 4 \\ + \left[12 - \Phi(12 + \gamma_1[Y_\tau]^2)\right] \alpha \\ + \left[13 - 30\Phi + 17\Phi^2 - 3\gamma_1[Y_\tau]^2\Phi(1-\Phi)\right] \alpha^2 \\ + \left[6 - 24\Phi + 30\Phi^2 - 12\Phi^3 - 3\gamma_1[Y_\tau]^2\Phi(1-\Phi)^2\right] \alpha^3 \\ + \left[1 - 6\Phi + 13\Phi^2 - 12\Phi^3 + 4\Phi^4 - \gamma_1[Y_\tau]^2\Phi(1-\Phi)^3\right] \alpha^4 = 0 \end{cases}}
$$

We can see that $\alpha$ is the root of a polynomial of order 4 which we solve numerically and choose the smallest real root as the solution. We reinject $\alpha$ into the first equation to find the parameter $\beta$.

##### 4th moment

The shape parameter of the Gamma distribution can also be inferred using the fourth-order normalized moment, known as kurtosis ($Kurt[Y_\tau]$). We use the same notation as before, and we start by explaining the kurtosis of $W_s$.

$$
Kurt[W_s] = E\left[\left(\frac{W_s - E[W_s]}{\sqrt{Var[W_s]}}\right)^4\right] = \frac{E[{W_s}^4] - 4E[W_s]E[{W_s}^3] + 6E[W_s]^2E[{W_s}^2] - 3E[W_s]^4}{Var[W_s]^2}
$$

To calculate $Kurt[Y_\tau]$ from $Kurt[W_s]$ we use the fourth-order normalized cumulant $\gamma_2$, also known as excess kurtosis, to use as before the cumulative property of cumulants on independent variables. $\gamma_2[Y_\tau] = Kurt[Y_\tau] - 3$, so we have :

$$
\gamma_2[Y_\tau] = Kurt[Y_\tau] - 3 = \frac{\sum_{s=0}^{S} Var[W_s]^2 \left(Kurt[W_s] - 3\right)}{\left(\sum_{s=0}^{S} Var[W_s]\right)^2}
$$

We now want to calculate the excess kurtosis for $Z$.
As before, using $E[Z^n] = \Phi\frac{\beta^n\Gamma(n+\alpha)}{\Gamma(\alpha)}$, we have :

$$
\begin{aligned}
\gamma_2[Z] &= E\left[\left(\frac{Z - E[Z]}{\sqrt{Var[Z]}}\right)^4\right] - 3 \\
&= \frac{E[Z^4] - 4E[Z]E[Z^3] + 6E[Z]^2E[Z^2] - 3E[Z]^4}{Var[Z]^2} - 3 \\
&= \frac{\Phi\beta^4(3+\alpha)(2+\alpha)(1+\alpha)\alpha - 4(\Phi\alpha\beta)(\Phi\beta^3(2+\alpha)(1+\alpha)\alpha) + 6(\Phi\alpha\beta)^2(\Phi\beta^2(1+\alpha)\alpha) - 3(\Phi\alpha\beta)^4}{(\Phi^2\alpha\beta^2 + \Phi(1-\Phi)\alpha^2\beta^2)^2} - 3 \\
&= \frac{6 + \alpha(11 - 8\Phi) + \alpha^2(6 - 12\Phi + 6\Phi^2) + \alpha^3(1 - 4\Phi + 6\Phi^2 - 3\Phi^3)}{\alpha\Phi + \alpha^2(1-\Phi)2\Phi + \alpha^3\Phi(1-\Phi)^2} - 3 \\
&= \frac{6 + \alpha(11 - 11\Phi) + \alpha^2(6 - 18\Phi + 12\Phi^2) + \alpha^3(1 - 7\Phi + 12\Phi^2 - 6\Phi^3)}{\alpha\Phi + \alpha^2(1-\Phi)2\Phi + \alpha^3\Phi(1-\Phi)^2}
\end{aligned}
$$

This gives us the following system:

$$\begin{cases} E[Y_\tau] = E[Z] \\ \gamma_2[Y_\tau] = \gamma_2[Z] \end{cases} \iff \begin{cases} E[Y_\tau] = \Phi\alpha\beta \\ \gamma_2[Y_\tau] = \frac{6+\alpha(11-11\Phi)+\alpha^2(6-18\Phi+12\Phi^2)+\alpha^3(1-7\Phi+12\Phi^2-6\Phi^3)}{\alpha\Phi+\alpha^2(1-\Phi)2\Phi+\alpha^3\Phi(1-\Phi)^2} \end{cases}$$

$$\iff \boxed{\begin{cases} \beta = \frac{E[Y_\tau]}{\Phi\alpha} \\ 6 \\ + [11 - \Phi(11 + \gamma_2[Y_\tau])]\,\alpha \\ + \left[6 - \Phi(18 + 2\gamma_2[Y_\tau]) + \Phi^2(12 + 2\gamma_2[Y_\tau])\right]\alpha^2 \\ + \left[1 - \Phi(7 + \gamma_2[Y_\tau]) + \Phi^2(12 + 2\gamma_2[Y_\tau]) - \Phi^3(6 + \gamma_2[Y_\tau])\right]\alpha^3 = 0 \end{cases}}$$

We can see that $\alpha$ is the root of a polynomial of order 3 which we also solve numerically and choose the smallest real root as the solution. As before, we reinject $\alpha$ into the first equation to find the parameter $\beta$.

#### Notes on the choice of the moments

In some cases, depending on the combination law $Y$, we may encounter situations where the calculated moment is negative, or, in the case of truncation, where we cannot find a positive real root to our $\alpha$ polynomial. These instances suggest that the moment does not exist. We have not pursued theoretical guarantees on the choice of moment based on the stratified contingency table data, but we can offer some empirical advice based on simulation and Supplementary Figure 1, 2, 3 and 4.

It appears that the choice of moment to use depends on the tail distribution of the combination law and the level of significance reached. Here are some guidelines:

- 2nd Moment : If the tail distribution of the combination law is not very spread out, it is preferable to use the 2nd moment, as higher-order moments may provide poorer estimates. This is a conservative choice that generally works well in practice.
- Higher Moments (3rd or 4th) : If the tail distribution is fairly spread out, higher-order moments (3rd or 4th) may yield better approximations. This is because they capture more information about the tail behavior of the distribution.

The Supplementary Figure **1** illustrates this phenomenon on the combination of 5 $p$-values from the same 5 strata of fixed size $N_s = 250$ with marginal $K_s = n_s = 24$. We can see that the 2nd moment is a good choice for the under-association case without truncation, while the 4th moment is more appropriate for the over-association case with truncation $\tau = 0.2$.

More generally, we have done several simulations to compare the choice of moment through the control of type I error at some predefined $\alpha$-level in function of sample size $N_s$, the marginals $n_s$ and $K_s$ and the value of the truncation $\tau$. When the set of combinations is less than 5 million, the type I error rate is calculated analytically from the exact distribution (we find where the inverse of the gamma distribution survival function for an alpha level lies in the exact distribution), so we can reach any alpha level. Alternatively, we run a simulation of 10000 combination samples and look at the type I error rate (we can then go to an alpha level of 0.0001). The results are shown in Supplementary Figure **3** and **4** with a zoomed case Figure **2**.

It is important to note that the spread of the tail of the distribution of the combination law is influenced by the maximum level of significance achievable (i.e., the smallest combined $p$-value possible), that is also influenced by the truncation. This means that in cases where extremely low $p$-values are achievable, the tail distribution may be more spread out, potentially warranting the use of higher moments. One way to study the spread of the tail of the distribution is to look at the ratio $\frac{Var[Y_\tau]}{\max(supp(Y_\tau))}$. If this ratio is less than 0.1, the tail distribution is more spread out, and higher moments may be more appropriate.

By default, we propose using the truncation threshold $\tau = 0.2$ to set aside strata with no association signal. With this value of $\tau$, we recommend to use 4th order moment to ensure the best approximation of the distribution. However, if the truncation threshold is set above, we recommend using the 2nd order.

In summary, users need to take into account the distribution characteristics of their specific data when deciding to use higher-order moments. Based on the above criteria, the choice of moment is also defined automatically. This empirical approach balances computational feasibility and approximation accuracy.

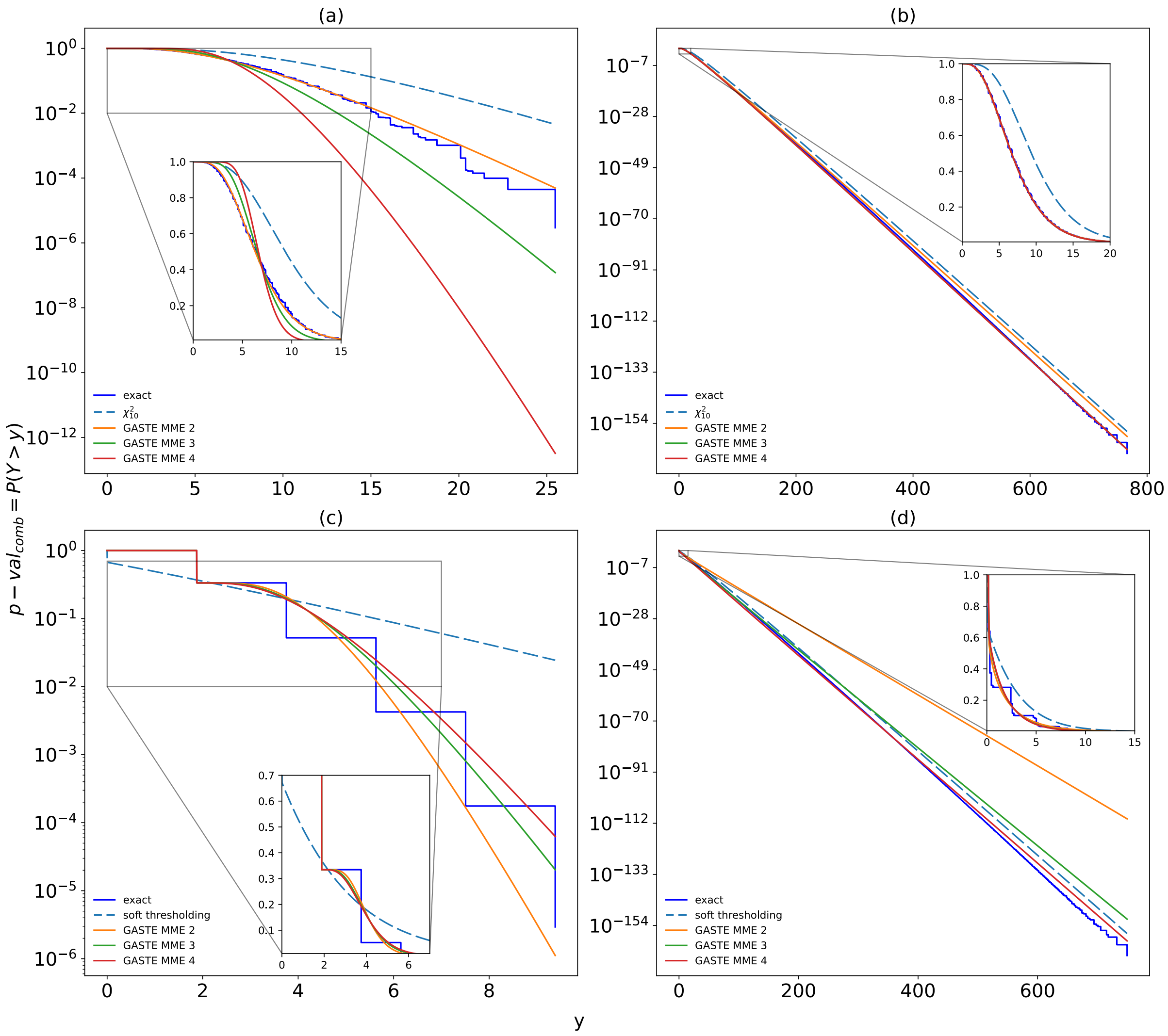


**Supplementary Figure 1**: Combination of 5 $p$-values from the same 5 strata of fixed size $N_s = 250$ with marginal $K_s = n_s = 24$. $(a)$ and $(b)$ are respectively the study of under-association and over-association without truncation and, $(c)$ and $(d)$ are respectively the study of under-association and over-association with truncation $\tau = 0.2$.

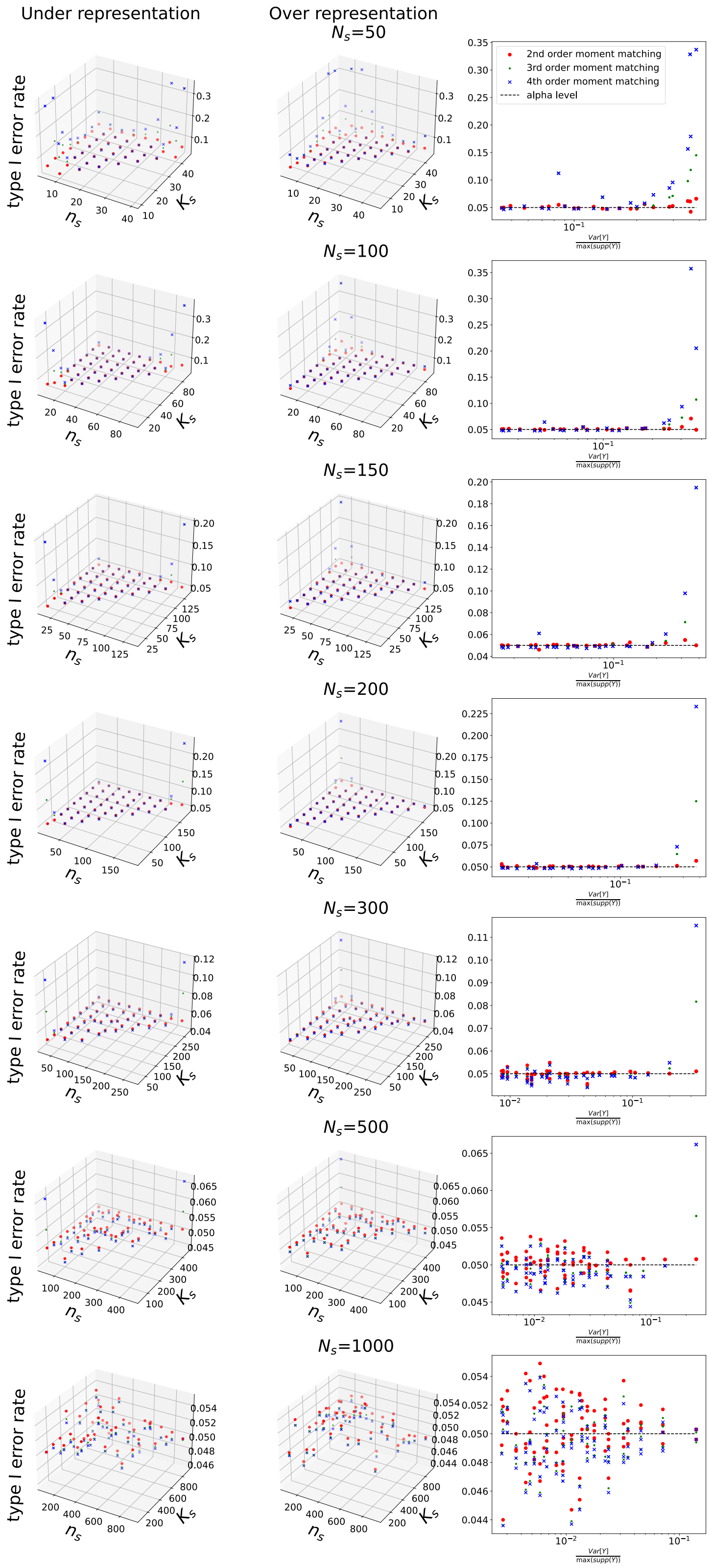


**Supplementary Figure 2**: Simulation of four strata contingency table for sample size $N_s = (50, 100, 150, 200, 300, 500, 1000)$ and marginals $K_s$ and $n_s$ taken 10% to 90% of $N_s$ with step of 10% and truncation $\tau = 1$. Study of type I error rate at $\alpha = 0.05$. The left subplot is the study of under-association, the center one is the study of over-association and the right subplot is the type I error rate in function of the spread of the tail of the distribution $(\frac{Var[Y_\tau]}{\max(supp(Y_\tau))})$. The gamma approximation using moment of order 2, 3 and 4 is compared.

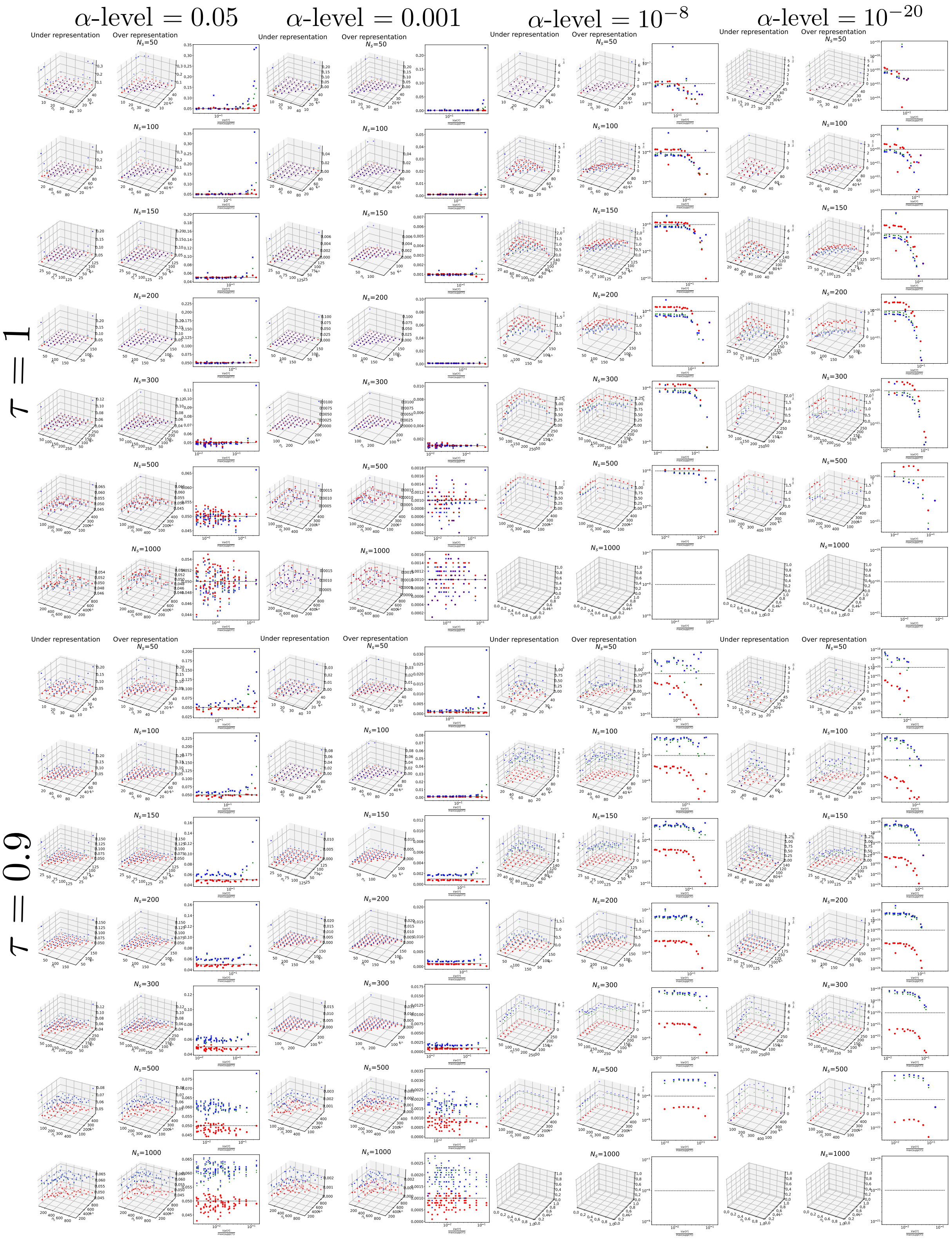


**Supplementary Figure 3**: Simulation of four strata contingency table for sample size $N_s = (50, 100, 150, 200, 300, 500, 1000)$ and marginals $K_s$ and $n_s$ taken 10% to 90% of $N_s$ with step of 10% and truncation $\tau = 1$ and $\tau = 0.9$. Study of type I error rate at $\alpha = (0.05, 0.001, 10^{-8}, 10^{-20})$ levels for under and over representation (same configuration of Figure 2).

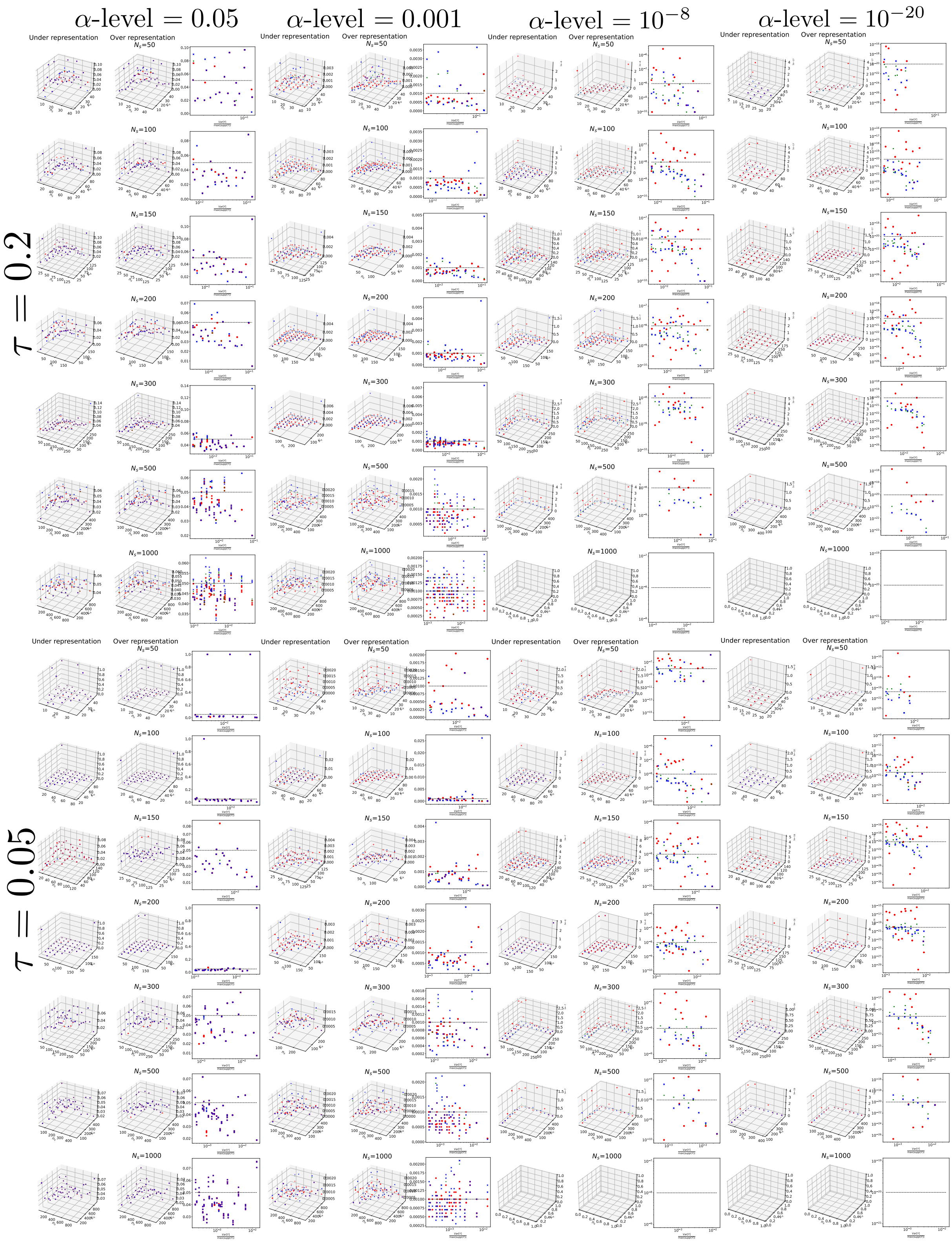


**Supplementary Figure 4**: Simulation of four strata contingency table for sample size $N_s = (50, 100, 150, 200, 300, 500, 1000)$ and marginals $K_s$ and $n_s$ taken 10% to 90% of $N_s$ with step of 10% and truncation $\tau = 0.2$ and $\tau = 0.05$. Study of type I error rate at $\alpha = (0.05, 0.001, 10^{-8}, 10^{-20})$ levels for under and over representation (same configuration of Figure 2).

### Material B : Numerical issue of exact calculation of the combined $p$-value

The expression of the combined $p$-values detailed at the core of the article is mathematically rigorous, but its practical implementation faces significant challenges. Due to the limitations of floating-point numbers, typically constrained to 64 bits, $p$-values are rounded to 0.0 if they fall below $5.10^{-324}$ and to 1.0 after 16 digits 9 after the decimal point. This rounding makes it possible to obtain identical $y$ combination values with different probabilities of realization. Additionally, rounding issues in the calculation of the $y$ combination value further complicate matters.

Supplementary Figure **5** illustrates three different ways to retain the probability of realization for a given $y$ combination:

- Minimum Probability (Blue) : For the same value of $y$, the minimum probability is used. This approach, however, leads to multiple identical $y$ combination values for different realization probabilities, due to the rounding issues.
- Maximum Probability (Green) : To be conservative, the maximum probability is used. This method also encounters similar issues with extremely close combination values, often within $10^{-14}$, which could either be due to rounding errors in the combination calculation or combinations that are genuinely very close in value.
- Rounding Strategy (Fuchsia) : The combination values are rounded to 10 powers minus the number of decimal digits plus three. This strategy ensures good conservatism and allows for a unique realization to be associated with a unique combination value.

The third approach appears to be the most practical and reliable, as it mitigates the issues arising from floating-point precision limitations and rounding errors, ensuring that each combination value corresponds to the maximum probability of realization.

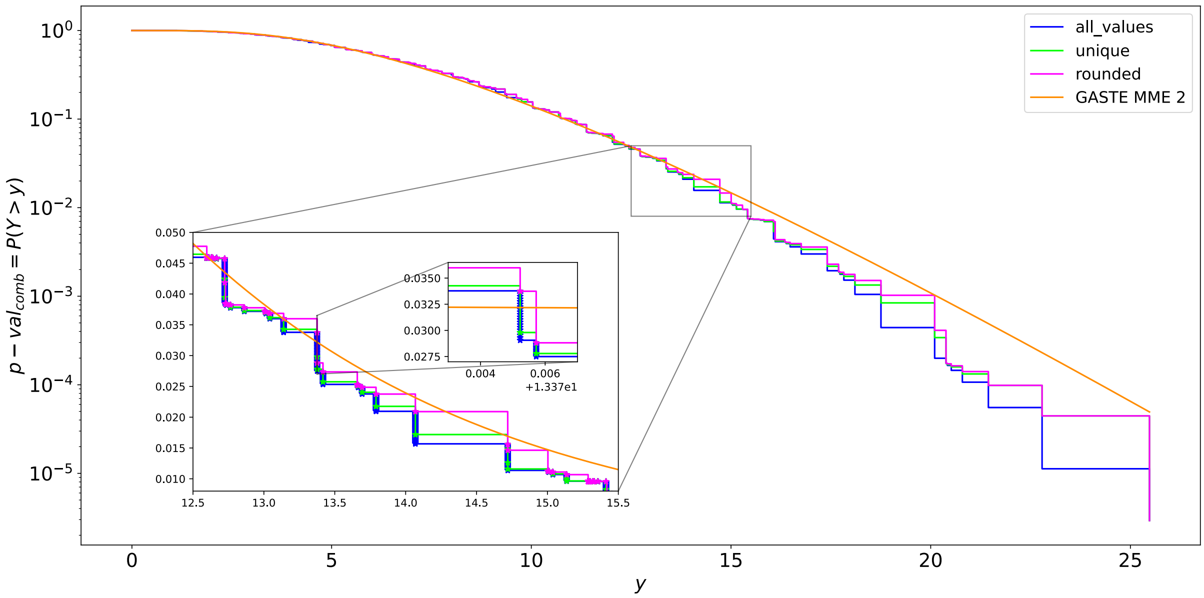


**Supplementary Figure 5**: Combination of 5 $p$-values from the same 5 strata of fixed size $N_s = 250$ with marginal $K_s = n_s = 24$ without truncation ($\tau = 1$), and three different ways to retain the probability of realization for a given $y$ combination

#### Material C : Time-accuracy simulation 1

In the Supplementary Figure **6**, on $x$ axis, we present the time of computation of the three method compared in Figure 1 of the paper and on $y$ axis a boxplot of the relative error of $-\log_{10}(p\text{-val}_{comb})$ of $\Gamma$ approximation and $\chi^2$ method compare to the exact law of combination $Y_1$ on all taken value. We can see that the $\Gamma$ approximation is very fast compared to the exact test (more than three order of magnitude lower) and very accurate compared $\chi^2$ distribution. The $p$-value obtained by $\chi^2$ distribution is far from the exact $p$-value, more than 40% of error on the magnitude of the $p$-value.

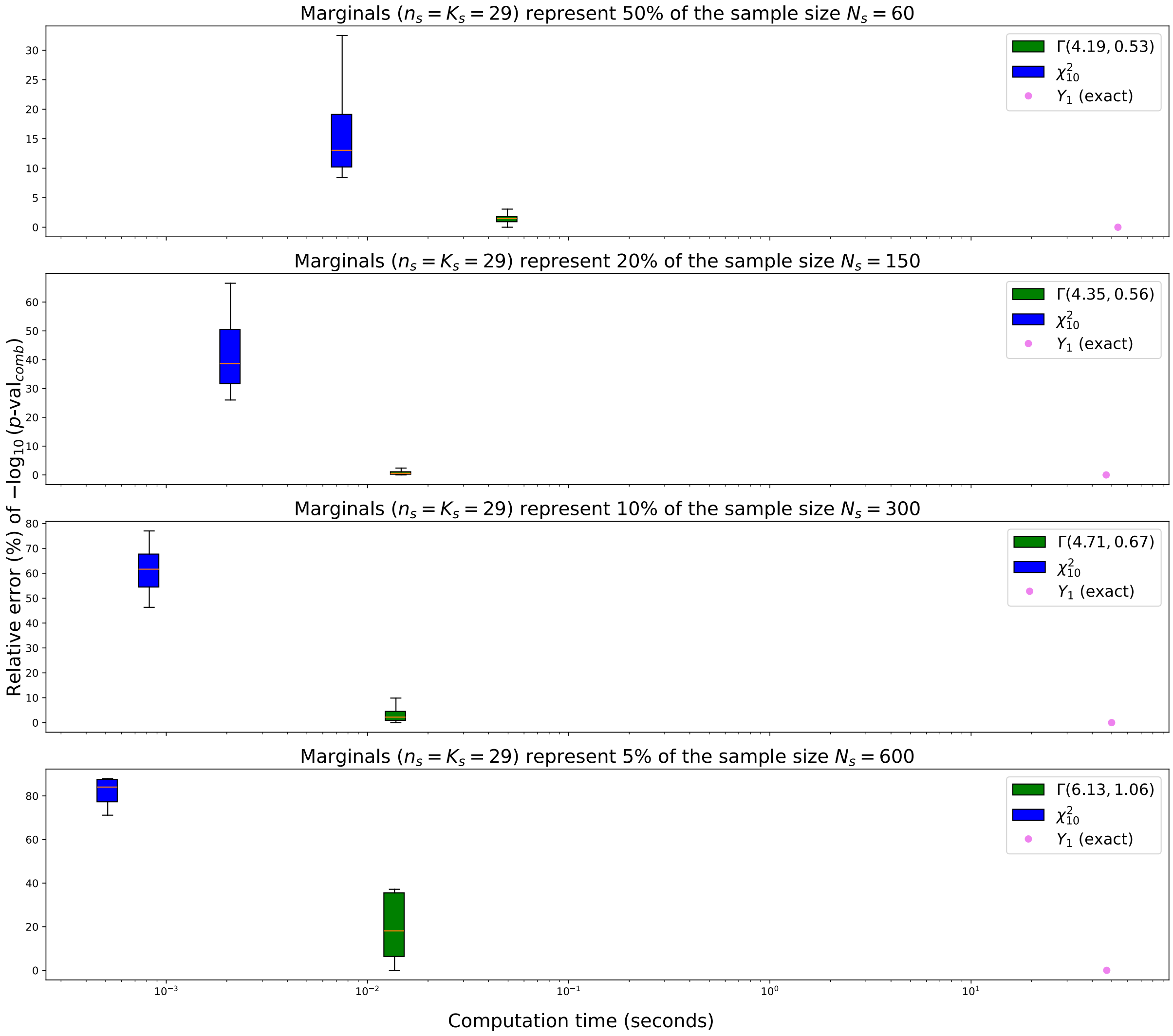


**Supplementary Figure 6**: Boxplot of relative error of the $\Gamma$ and $\chi^2$ distribution compare to the exact distribution of the combined $p$-value in the settings of Figure 1 of the paper

#### Material D : Figure time complexity

Here we present the computational time complexity of the exact distribution and the Gamma approximation as a function of the number of strata. The computational time complexity of the exact distribution is exponential with respect to the number of strata. This exponential growth in complexity is due to the increasing number of possible configurations of the contingency tables that must be considered as the number of strata increases.

In contrast, the computational time complexity of the Gamma approximation is linear with respect to the number of strata. This linear complexity arises because the Gamma approximation leverages the moments of the distribution, which can be computed in a straightforward manner without enumerating all possible configurations. As a result, the Gamma approximation scales much more efficiently as the number of strata increases.

The Supplementary Figure **7** illustrates the comparison of computation time for exact and Gamma approximation calculation as a function of the number of strata, between 2 and 10 for support ($HG(15, 5, 5)$). The support is tiny to be able to compute several strate.

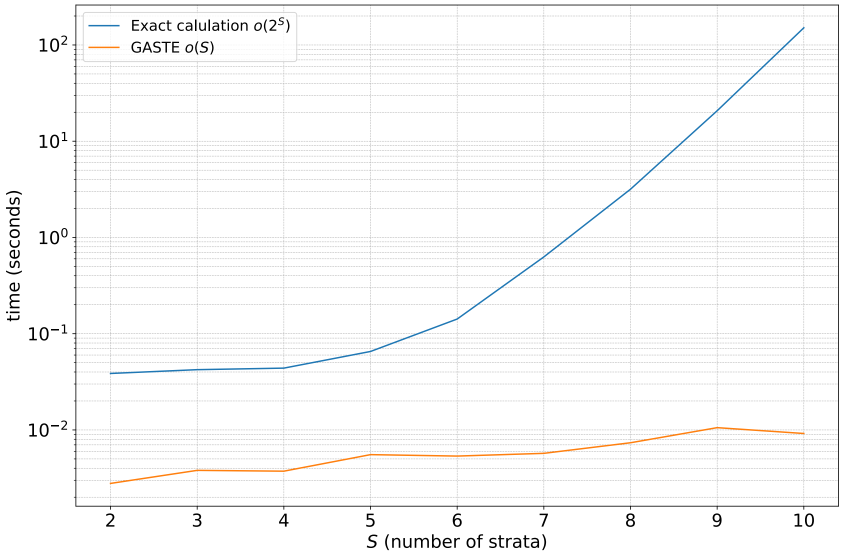


**Supplementary Figure 7**: Comparison of computation time for exact and Gamma approximation calculation in function of the number of strata for a tiny support ($HG(15, 5, 5)$)

### Material E : Example of Forest plot on Orchamp data

#### E.1 Opposite significant coexistence

We present a forest plot stratified by different environment of the coexistence between plants species *Agrostis schraderiana* and *Rumex arifolius* on the Orchamp data (Supplementary Figure **8**). The $p$-values of overall effect are calculated using the Gamma approximation method. We can see that the combined $p$-value is significant for the under-association and over-association as the same time. This means that the coexistence between these two species is significant in different way in the different environments. Through the forest plot, the environment *BOU_1620* brings a significant level of the overall over-association, which is a closed environment like forest. On the other hand, the environments *ANT_1750* bring a significant level of the overall under-association, which is an open environment like grassland. This result is not inconsitent with the ecological knowledge, that a harsh environment leads plants to evolve in the same restricted area, and inversely, a rich environment leads plants to expand and conquer an area, producing competition between plants. Here, for this pair, their presence in the forest seems to correspond to a harsher environment (e.g. environmental constraints linked to sunshine), whereas in the grassland, they seem to be more in competition due to a lesser environmental constraint.

Additionally, we can note that due to the opposite effect, the overall effect given by CMH is completely no significant ($p$-value=0.79).

| | Is plant A: Is plant B | Is plant A: Is not plant B | Is not plant A: Is plant B | Is not plant A: Is not plant B | $p_s^-$ | $p_s^+$ | Log Odd ratio with 95% CI | %W(fixed) |
|---|---|---|---|---|---|---|---|---|
| BOU_1620 | 26 | 87 | 3 | 180 | 1.0000 | 2.18e-09 | 2.887 [1.664, 4.109] | 3.6% |
| ANT_1750 | 6 | 51 | 109 | 134 | 3.32e-07 | 1.0000 | -1.934 [-2.816, -1.051] | 75.8% |
| TEN_1890 | 0 | 4 | 1 | 292 | 0.9865 | 1.0000 | 3.597 [0.062, 7.133] | 0.1% |
| PAI_1900 | 8 | 44 | 32 | 216 | 0.7645 | 0.3868 | 0.205 [-0.635, 1.045] | 19.2% |
| BON_1900 | 0 | 12 | 1 | 286 | 0.9599 | 1.0000 | 2.478 [-0.966, 5.921] | 0.2% |
| MER_2000 | 0 | 1 | 6 | 287 | 0.9796 | 1.0000 | 3.175 [-0.315, 6.664] | 0.1% |
| TAN_2100 | 2 | 78 | 1 | 203 | 0.9783 | 0.1924 | 1.650 [-0.765, 4.064] | 1.1% |
| FE Model **Overall effect CMH**, test of $\log(\theta)=0$, CMH=0.0660 **p = 0.7973** | | | | | | | 0.104 [-0.286,0.495] | 100.0% |

Test for **Homogeneity**, test of $\theta_i=\theta_j$, BD=60.5897 **p = 3.42e − 11**

**Overall effect, Stratified Exact test (GASTE)**, $\#supp(Y) = 2.28e+06$

test of $\forall s$, $\log(\theta_s) \geq 0$, $Y_1^-$ =30.5694, $\mathbf{p^-}$ **= 1.06e − 05**

test of $\forall s$, $\log(\theta_s) \leq 0$, $Y_1^+$ =45.0797, $\mathbf{p^+}$ **= 1.51e − 07**

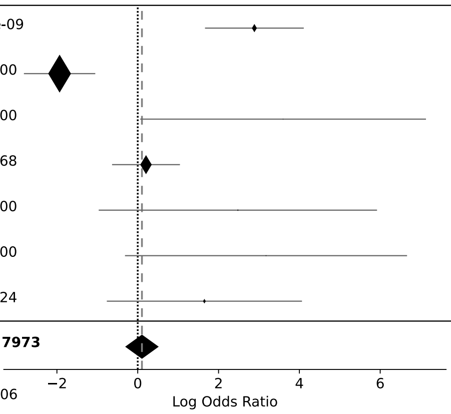


**Supplementary Figure 8**: Forest plot of the combined $p$-value of the under-association and over-association of the coexistence between plants species *Agrostis schraderiana* and *Rumex arifolius*

| | Is plant A: Is plant B | Is plant A: Is not plant B | Is not plant A: Is plant B | Is not plant A: Is not plant B | $p_s^-$ | $p_s^+$ | Log Odd ratio with 95% CI | %W(fixed) |
|---|---|---|---|---|---|---|---|---|
| SAP_1870 | 18 | 43 | 51 | 174 | 0.8976 | 0.1733 | 0.356 [-0.276, 0.989] | 8.5% |
| RBI_1950 | 4 | 4 | 162 | 128 | 0.5070 | 0.7558 | -0.236 [-1.641, 1.170] | 2.4% |
| MON_1970 | 2 | 45 | 38 | 209 | 0.0263 | 0.9946 | -1.409 [-2.867, 0.049] | 6.5% |
| CHAM_2020 | 5 | 119 | 11 | 146 | 0.2107 | 0.9098 | -0.584 [-1.668, 0.501] | 5.2% |
| BON_2140 | 0 | 26 | 61 | 210 | 0.0019 | 1.0000 | -2.715 [-5.528, 0.098] | 5.9% |
| RBI_2150 | 0 | 13 | 42 | 225 | 0.1149 | 1.0000 | -1.580 [-4.423, 1.264] | 2.2% |
| CHAM_2180 | 6 | 39 | 22 | 147 | 0.6317 | 0.5624 | 0.028 [-0.942, 0.997] | 4.4% |
| RIS_2195 | 14 | 55 | 55 | 167 | 0.2766 | 0.8225 | -0.258 [-0.919, 0.404] | 11.5% |
| NEV_2200 | 5 | 102 | 11 | 178 | 0.4488 | 0.7493 | -0.232 [-1.316, 0.853] | 4.2% |
| DEV_2340 | 10 | 10 | 58 | 211 | 0.9984 | 0.0068 | 1.291 [0.368, 2.215] | 2.2% |
| MOU_2355 | 6 | 26 | 81 | 186 | 0.1215 | 0.9468 | -0.635 [-1.560, 0.290] | 7.8% |
| VAL_2450 | 3 | 71 | 23 | 197 | 0.0682 | 0.9797 | -1.016 [-2.250, 0.217] | 6.2% |
| MOU_2470 | 6 | 51 | 35 | 162 | 0.1332 | 0.9401 | -0.608 [-1.529, 0.314] | 7.8% |
| VAN_2630 | 31 | 61 | 94 | 86 | 0.0026 | 0.9989 | -0.766 [-1.288, -0.244] | 23.4% |
| VAL_2710 | 1 | 3 | 124 | 105 | 0.2580 | 0.9552 | -1.265 [-3.543, 1.013] | 1.8% |
| FE Model **Overall effect CMH**, test of $\log(\theta)=0$, CMH=12.4870 **p = 4.10e − 04** | | | | | | | -0.411 [-0.649,-0.174] | 100.0% |

Test for **Homogeneity**, test of $\theta_i=\theta_j$, BD=35.1057 **p = 0.0014**

**Overall effect, Stratified Exact test (GASTE)**, $\#supp(Y) = 1.85e+21$

test of $\forall s$, $\log(\theta_s) \geq 0$, $Y_1^-$ =62.1460, $\mathbf{p^-}$ **= 9.35e − 06**

test of $\forall s$, $\log(\theta_s) \leq 0$, $Y_1^+$ =16.7236, $\mathbf{p^+}$ **= 0.8616**

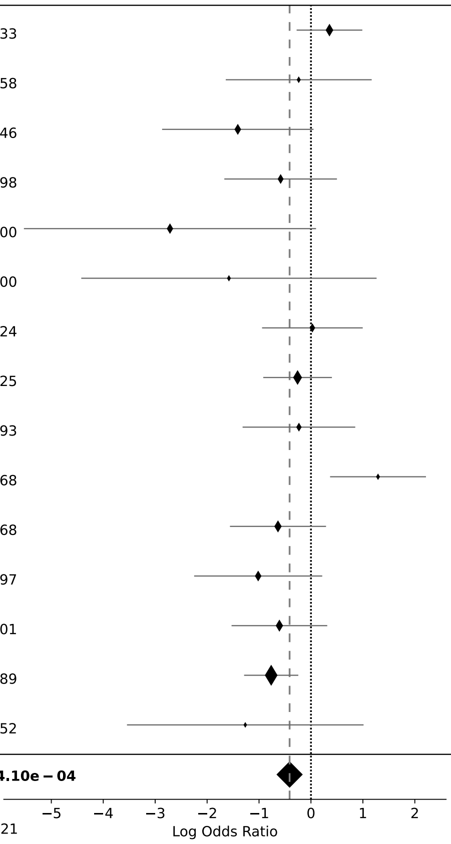


**Supplementary Figure 9**: Forest plot of the combined $p$-value of the under-association and over-association of the coexistence between plants species *Carex sempervirens subsp. sempervirens* and *Festuca violacea*

#### E.2 Widespread significant

In this example (Supplementary Figure **9**), unlike the previous example, the overall significance signal is not carried strongly by a single plot, but is much more widespread across all plots with lower local significance. This can be interpreted as a more generalized signal of coexistence between the two plant species *Carex sempervirens subsp. sempervirens* and *Festuca violacea* across all plots. We can see that the GASTE method also detects this generalized coexistence signal very well, and more significantly than the CMH method.

### Material F : Asymptotic convergence of Gamma model

We can also ask about the relevance of the choice of the Gamma model to approximate the exact distribution. We can try to see if the Gamma approximation converges asymptotically to a $\chi^2$ when the $p$-values approach a continuous uniform distribution. To do this, we can vary the size $N_s$ of the hypergeometric distribution. Indeed, when the size $N_s$ is large and the marginals $K_s$ and $n_s$ are of the same order of magnitude, the support of the hypergeometric distribution is large and there are therefore as many possible $p$-values. The $p$-values can therefore possibly be better distributed and approach a uniform distribution.

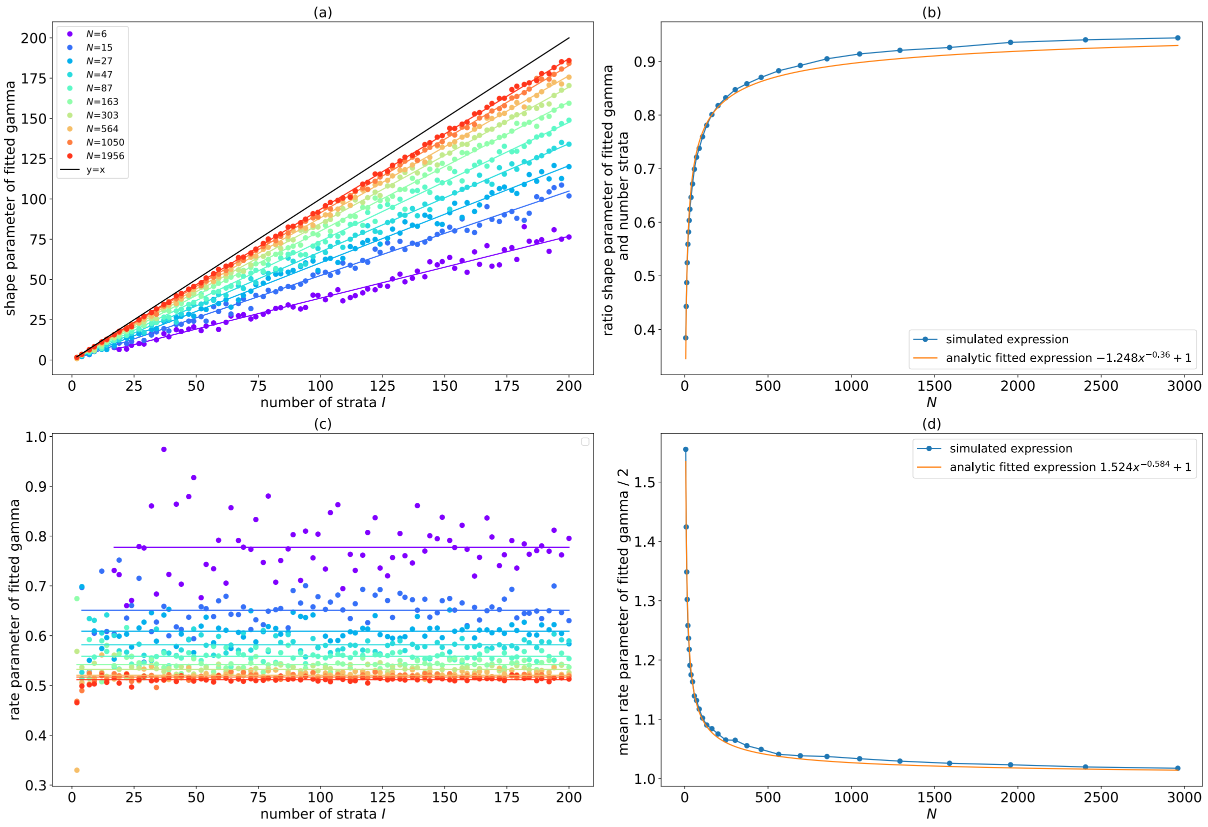


**Supplementary Figure 10**: Simulation of different strata with all same sample size $N$ (call now here $N$) and margins draw randomly from 1 to $N$. For each case, we simulate the untruncated combination distribution and approximate it by a classic Gamma distribution. ($a$) shape parameter in function of number of strata $S$ for different sample $N$, ($b$) same thing for rate parameter, ($c$) ratio of shape parameter and number of strata $S$ in function of sample size, ($d$) mean value of rate parameter in function of sample size $N$.

We recall that here $Y_1 = -2\sum_{s=1}^{I}\log(P_s)$ where $P_s$ follow a sub-uniform distribution from hypergeometric test. The following model is assumed, $Y_1 \approx Gamma(I\,g(N), \frac{h(N)}{2})$, with

$\lim_{x \to \infty} g(x) = 1$ and $\lim_{x \to \infty} h(x) = 1$. We found by simultation $g(x) = -1.248x^{-0.36} + 1$ and $h(x) = 1.524x^{-0.584} + 1$. As we can see on sub-figure $(b)$ and $(d)$ of the Supplementary Figure **10** the model is very well-fitted to the simulation.

We can see that the shape parameter $\alpha$ and the rate parameter $\beta$ of the Gamma distribution converge respectively to the number of strata $S$ and 0.5 when the sample size $N_s$ increases. That is the parameters of Gamma distribution corresponding to a $\chi^2_{2S}$. This is consistent with the fact that the $p$-values are better distributed and approach a uniform distribution when the sample size increases. So our *Gamma* model seems to be a good approximation of the exact distribution of the combined $p$-value and generalizes the $\chi^2$ distribution of the combined $p$-value.
